# Deep Diffusion MRI Registration (DDMReg): A Deep Learning Method for Diffusion MRI Registration

**DOI:** 10.1101/2021.03.04.433968

**Authors:** Fan Zhang, William M. Wells, Lauren J. O’Donnell

## Abstract

In this paper, we present a deep learning method, DDMReg, for accurate registration between diffusion MRI (dMRI) datasets. In dMRI registration, the goal is to spatially align brain anatomical structures while ensuring that local fiber orientations remain consistent with the underlying white matter fiber tract anatomy. DDMReg is a novel method that uses joint whole-brain and tract-specific information for dMRI registration. Based on the successful VoxelMorph framework for image registration, we propose a novel registration architecture that leverages not only whole brain information but also tract-specific fiber orientation information. DDMReg is an unsupervised method for deformable registration between pairs of dMRI datasets: it does not require nonlinearly pre-registered training data or the corresponding deformation fields as ground truth. We perform comparisons with four state-of-the-art registration methods on multiple independently acquired datasets from different populations (including teenagers, young and elderly adults) and different imaging protocols and scanners. We evaluate the registration performance by assessing the ability to align anatomically corresponding brain structures and ensure fiber spatial agreement between different subjects after registration. Experimental results show that DDMReg obtains significantly improved registration performance compared to the state-of-the-art methods. Importantly, we demonstrate successful generalization of DDMReg to dMRI data from different populations with varying ages and acquired using different acquisition protocols and different scanners.

## 1 Introduction

Diffusion magnetic resonance imaging (dMRI) is an advanced neuroimaging technique that measures the random diffusion of water molecules in the brain [1]. dMRI data includes multi-dimensional, orientation-dependent signals that describe not only the strength but also the orientation of water diffusion in brain tissues. As a result, dMRI provides a unique technique to enable in-vivo fiber tracking (tractography) of white matter fiber tracts [2] and to estimate the underlying cellular microstructure of brain tissues [3]. Registration of dMRI data is a crucial step in applications such as population-based analyses of diffusion MRI data in health and disease [4], [5] and construction of brain anatomical atlases [6]–[8].

dMRI registration is a challenging task. In traditional neuroimaging data (e.g., T1- or T2-weighted MRI), the images are scalar-valued volumes, representing anatomically meaningful numeric intensity values that are specific to certain brain structures. Registration is performed usually by minimizing the intensity dissimilarity between the volumes to spatially align the corresponding anatomical structures in the brain, such as white matter (WM), gray matter (GM) and cerebrospinal fluid (CSF) as well as their subdivisions [9]. dMRI data, however, describe not only the strength but also the orientation of water diffusion. The orientation information encoded in dMRI data is important to reveal the underlying fiber orientation in the white matter. Therefore, dMRI registration should not only spatially align anatomical structures of the entire brain, but also ensure that local fiber orientations remain consistent with the underlying white matter anatomy after image transformations [10]–[14].

Many methods have been proposed for dMRI registration, which can be generally categorized based on the input data to the algorithms. The most straightforward approaches use representative scalar images derived from the dMRI data, so that existing methods developed for traditional imaging data can directly use or easily be adapted for dMRI registration [5], [15]. Currently, the fractional anisotropy (FA) image [3] is the most popular and has been widely used in research studies [5], [15]–[20]. Nevertheless, despite the wide usage of scalar images for dMRI registration, this approach does not use water diffusion orientation information contained in the dMRI data and thus discards important information to ensure fiber tract consistency.

A number of approaches for dMRI registration have used input data containing diffusion orientation information. Early work used diffusion tensor imaging (DTI) [11], [21]–[26], where each voxel is a 2nd-order Cartesian tensor that can estimate the principal water diffusion direction. However, a known issue is that DTI can not adequately model more complex fiber configurations such as crossing fibers that are prevalent in much of the brain [27]. Consequently, many dMRI registration methods have used more advanced diffusion models that can estimate multiple fiber orientations at each voxel. The widely used models include higher-order diffusion tensor models [28], diffusivity functions [29], Gaussian mixture fields [30], diffusion orientation distribution functions (ODF) [9], [31], and fiber orientation distributions (FOD) [32]–[35]. ODF and FOD models are often represented using spherical harmonic bases to facilitate registration [9], [31]–[35]. Registration based on these higher-order models has shown to better align brain regions in the presence of crossing fibers. In particular, methods based on the FOD are popular and software packages are available [36] to enable clinical research [37]–[39]. However, because the larger-scale fiber tract geometry underlying the local model is unknown [40], the registration solution may be ambiguous, especially in the presence of crossing tracts of similar amplitude.

Finally, a number of approaches for dMRI registration have used input data representing white matter fiber tracts. Several methods have been proposed to perform fiber tractography and then register based on fiber streamline trajectory information [41]–[45]. Other tractography-based registration methods register volumetric maps computed from the fiber streamlines, such as fiber tract probability (or density) maps [46], [47] and currents (maps computed based on the tangent vector of the streamline trajectory) [48]. These methods have shown good performance to align white matter tracts. Nevertheless, these methods are designed specifically for tract registration, without considering other brain structures.

In terms of methodology, almost all existing dMRI registration methods apply the traditional medical image registration formulation, where the algorithms solve an optimization problem to iteratively align an image pair based on an energy function [49]. Recent advances in deep learning techniques have shown significantly improved computational speed and registration accuracy in medical image registration tasks [50]–[52]. A handful of recent publications have shown the potential of registration of dMRI data using deep learning. One deep learning method demonstrated that the incorporation of local fiber orientation information (using diffusion tensor images) improved registration accuracy over registration based on the scalar T2-weighted image alone [53]. Another deep learning approach performed simultaneous segmentation and registration of diffusion MRI data, where the registration applied the popular approach of scalar FA image registration to align based on whole-brain tissue microstructure [20]. However, to our knowledge, existing deep learning methods for dMRI registration do not yet leverage both whole-brain and tract-specific information.

In this study, we propose a novel deep learning method that uses joint whole-brain and tract-specific information for registration of dMRI data, which we refer to as the *Deep Diffusion MRI Registration* (*DDMReg*) method. There are several contributions and advantages of our method. First, DDM-Reg is an unsupervised learning method for deformable registration between dMRI datasets. While DDMReg enables deformable registration to account for local nonlinear differences of the brains from different individuals, the training process requires only rigidly aligned images and does not require nonlinearly pre-registered training data or the corresponding deformation fields as ground truth. Second, DDMReg contains a novel multi-input fusion network architecture, where multiple U-net-based subnetworks (based on the successful VoxelMorph [54] framework) for registration are trained on different types of input data derived from dMRI and are subsequently fused for final deformation field estimation. In this way, DDMReg enables good registration of anatomical structures from the entire brain, while ensuring good fiber tract alignment with respect to the underlying white matter anatomy. Third, in order to handle the computational limit due to GPU memory, we propose a novel multi-deformation fusion subnetwork to leverage backbone networks, where weights of a model for registering a certain type of input data can be pretrained.

## 2 Methods

The goal of the proposed DDMReg method is to compute a deformation field (*ϕ*) to register a pair of dMRI datasets, i.e., registering a moving (*m*) to a target (*t*) dMRI dataset. Fig. 1 gives an overview of the method. From the raw dMRI data, also known as the diffusion weighted imaging (DWI) data, we first compute input data to our network (Sec. 2.1). Two types of input data are computed, in which one type (i.e., FA) is generally useful for aligning anatomical structures from the whole brain while the other type (i.e., tract orientation maps, TOMs, proposed in [55]) is useful for aligning fibers of specific anatomical white matter tracts. For each type of input data, a U-net-based registration subnetwork (the same network architecture proposed in [54]) is trained to compute a deformation field that is specific to the input data type. A multi-deformation fusion subnetwork is then used to combine all computed deformation fields to estimate the final deformation field. One of the benefits of the multi-deformation fusion subnetwork is that the different registration subnetworks constrain each other, while cooperating to optimize the entire network. Following that, a spatial transformation subnetwork [56] and a fiber reorientation subnetwork are used to warp the moving input data into the target subject space (Sec. 2.2). Given the large number of registration subnetworks (a total of 41 in our study) that is difficult to fit in the GPU at once, we pretrain model weights of the registration subnetworks that are specific for the white matter fiber tracts, and we use them as backbones in our overall network (Sec. 2.4).

**Fig. 1.**
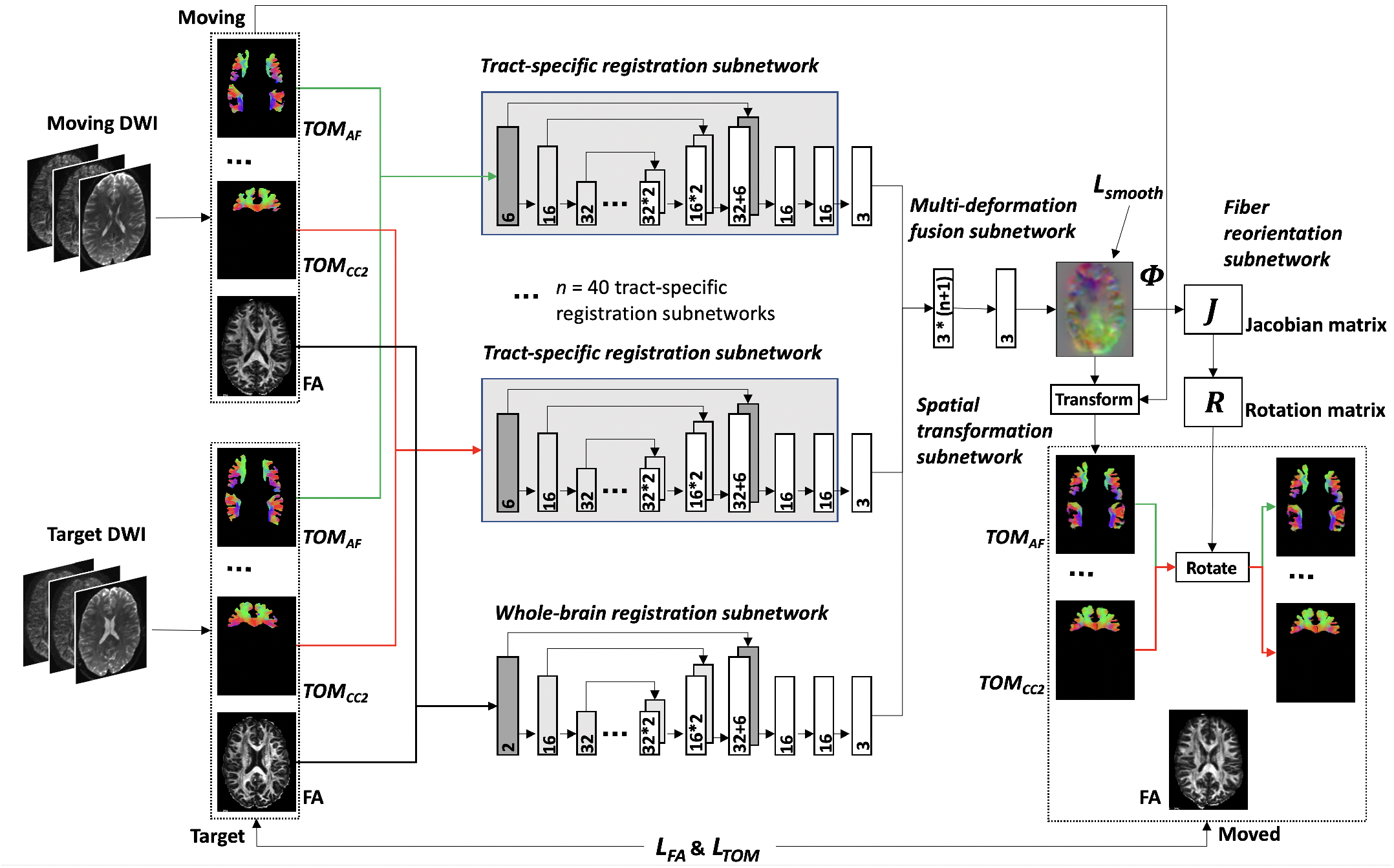
Method overview of DDMReg. From DWI data of the moving and target subjects, FA and 40 TOM images are computed as input to our network. Whole-brain (n = 1) and tract-specific (n = 40) registration subnetworks are trained to compute deformation fields specific to the input data (FA or TOM). A multi-deformation fusion subnetwork is used to combine all computed deformation fields to estimate the final deformation field. A spatial transformation subnetwork is used to warp the moving input data (i.e., FA and TOMs). A fiber reorientation subnetwork is used to reorient the warped TOM to ensure the local fiber orientation remains anatomically correct after warping. The model weights of the convolutional layers (gray rectangle) in each tract-specific registration subnetwork are pretrained to reduce computational burden (see Fig. 2).

### 2.1 Network Input

Input data for dMRI registration should be generally descriptive of the different types of anatomical structures in the brain so that it can be used to align the whole brain. Meanwhile, input data should also be useful for describing local fiber orientations to ensure alignment of individual white matter fiber tracts, in particular in the presence of crossing fibers. In our method, we propose to include these two kinds of input images.

For the description of the whole brain, we use the FA image. FA measures the water diffusion anisotropy of the tissues in the brain [3], one of the most important quantitative measures that dMRI can measure. The FA image gives a good contrast for anatomical brain structures such as WM, GM and CSF, and it has been widely used for dMRI registration tasks [5], [15]–[20]. (In Supplementary Fig. S1, we also test several other images that have been used for dMRI registration, including diffusion baseline images, diffusion tensor images and FOD peak images. We show that FA generates the best registration performance for our proposed framework.)

For the description of specific white matter tracts, we use tract orientation maps (TOMs) [55]. The TOM contains local fiber orientations of a specific white matter tract (e.g., arcuate fasciculus). At each voxel, a TOM contains one 3D vector (one peak), which represents the local mean fiber streamline orientation of a particular fiber tract. There are several advantages of applying TOMs in our framework. First, each TOM is tract-specific. In the presence of crossing fibers (where voxels contain multiple fiber orientation peaks) the TOM of a tract can identify the orientation peak that is specific to the tract. Second, the TOM is learned using CNNs based on fiber orientations estimated directly from fiber streamlines (see [55] for details). In this case, TOMs encode geometric information of fibers so that registering using TOMs can be helpful to ensure geometric alignment of fiber tracts. Third, computation of TOMs can be done using the GPU, which is time efficient. Fourth, TOMs are stored as volumetric images (a 3D volume with three channels representing the fiber orientation at each voxel), which can be easily used as input to CNNs.

### 2.2 Architecture

Our proposed network contains four subnetworks: 1) a whole-brain registration subnetwork, 2) *n* tract-specific registration subnetworks, 3) a multi-deformation fusion subnetwork, 4) a spatial transformation subnetwork, and 5) a fiber reorientation subnetwork.

The whole-brain registration subnetwork learns a deformation field *ϕ_wb_* between the target and moving FA images, i.e., FA(*t*) and FA(*m*). We use the Unet-based architecture proposed in VoxelMorph, a popular library for medical image registration [54] (see Fig. 1). Specifically, the subnetwork takes as input a 2-channel 3D volume formed by concatenating FA(*t*) and FA(*m*). 3D convolutions are performed in both the encoder and decoder stages using a kernel size of 3, and a stride of 2 (see [54] for other detailed parameter settings). The last layer of this subnetwork is a 3-filter 3D convolutional layer, which outputs a 3-channel 3D volume, i.e., the estimated deformation field *ϕ_wb_*.

Each tract-specific registration subnetwork learns a deformation field 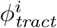 between the target and moving TOM images for a certain white matter tract *i,* i.e., TOM^*i*^(*t*) and TOM^*i*^(*m*). Here, *i* ∈ [1,…,*n*], where *n* is the total number of tract-specific registration subnetworks. We use a similar Unet-based architecture as applied in the whole-brain registration subnetwork, but the input is a 6-channel 3D volume formed by concatenating TOM^*i*^(*t*) and TOM^*i*^(*m*). The last layer is also a 3-filter 3D convolutional layer that outputs the estimated deformation field 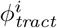. In our study, we have a total of *n* = 40 tract-specific subnetworks, corresponding to the number of tract-specific TOMs available in [55] (see Sec. 3 for details).

The multi-deformation fusion subnetwork combines the whole-brain deformation field and all tract-specific deformation fields to calculate the final deformation field *ϕ_final_*. The multi-deformation fusion subnetwork is based on our recently proposed dual-stream network to fuse deformation fields for multimodal image registration [57]. The idea is to concatenate different deformation fields, followed by a 3-filter 3D convolution to estimate the final deformation field. In our study, the multi-deformation fusion layer takes an input 3D volume with 3 × (*n* +1) channels formed by concatenating *ϕ_wb_* and all *n ϕ_tract_*, performs 3D convolution with size 3 × 3 × 3, and outputs a 3D volume with 3 channels, i.e., *ϕ_final_*.

The spatial transformation subnetwork warps the moving FA and TOM images into the target space using *ϕ_final_* for loss computation (see Sec. 2.3). This step is done using the spatial transformer network (STN) proposed in [56].

The fiber reorientation subnetwork reorients each warped TOM so that the local fiber orientation remains anatomically correct after warping. We apply the widely used finite strain (FS) strategy, which was designed for diffusion tensor reorientation in dMRI registration [10]. The FS strategy has been successfully applied for tensor reorientation in a recent deep learning image registration method [53]. Specifically, in our fiber reorientation subnetwork, the local Jacobian matrix (*J*) is computed at each voxel from the deformation field *ϕ_final_*, followed by a polar decomposition of *J* to obtain a unitary matrix *R* (a pure rotation matrix). Then, R is then used to reorient each TOM.

### 2.3 Loss Function

Our overall loss function *L_final_* consists of three components, including: 1) *L_FA_* that penalizes differences of FA to enforce whole brain registration accuracy, 2) 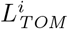 that penalizes differences of each TOM^*i*^ to enforce white matter tract registration accuracy, 3) *L_smooth_* that penalizes local spatial variations in the deformation fields to encourage deformation smoothness:

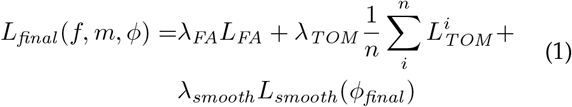

where *λ*_FA_, *λ_TOM_* and *λ_smooth_* are the weighting parameters that denote the relative importance of the three components to balance the overall registration accuracy and the smoothness of the deformation field (see Sec. 2.5 for detailed settings). For *L_FA_* and 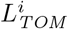, we use the widely applied mean squared voxelwise difference, i.e., the mean squared error (MSE) between the moved and target images, that has been shown to be effective for deep learning image registration in VoxelMorph [54]. (In Supplementary Fig. S1, we also test additional difference measures, including normalized cross correlation (NCC) [58] for FA and cosine similarity for TOM [55], where we find that MSE obtains the best performance.) For *L_smooth_*, we use an L2-norm of the spatial gradients of the deformation field, as suggested in [54].

### 2.4 Pretraining Tract-Specific Backbone Subnetworks

Leveraging pretrained network models as backbones has shown to be successful for improving both effectiveness and efficiency in medical image computing tasks [59]–[61]. In our work, given the large number of registration subnetworks and the limit of GPU memory, it is difficult to train all U-net models at the same time. To handle this, we propose to pretrain the U-net model for each tract-specific registration subnetwork (as illustrated in Fig. 2). Specifically, given each tract-specific TOM^*i*^, we pretrain the corresponding U-net model with a loss function based on only the TOM data:

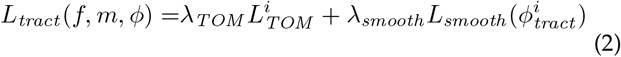

where 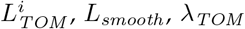 and *λ_smooth_* are the same as in Eq. (1). These pretrained U-net models are then used as backbones to compute tract-specific deformation fields in our overall network.

**Fig. 2.**
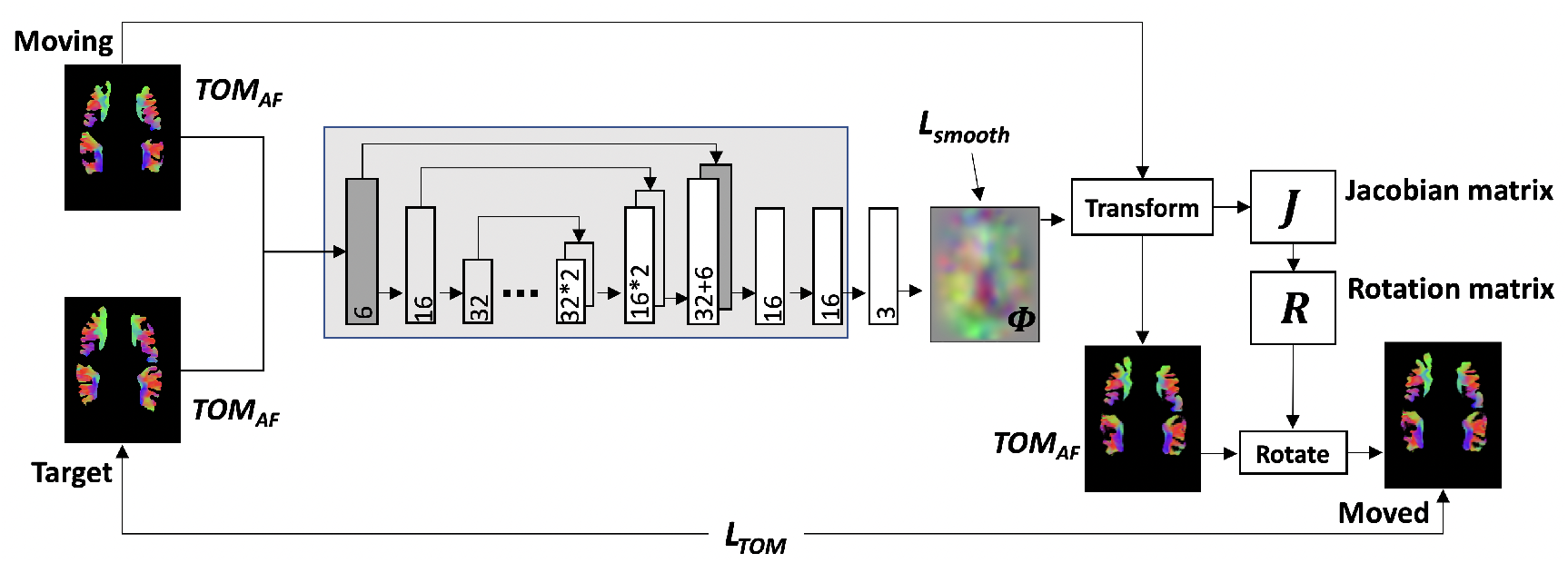
Pretraining tract-specific backbone subnetwork. After pretraining, the weights of the convolutional layers (in gray rectangle) are directly used in the overall network (Fig. 1) as a backbone to compute the tract-specific deformation field.

For training of the overall network, the model weights of the pretrained tract-specific registration subnetworks (the convolutional layers indicated using the gray rectangles in Figs. 1 and 2) are locked during backpropagation. However, the weights of the whole-brain registration subnetwork are not pretrained, and are updated during backpropagation. In this way, the whole-brain registration subnetwork training can benefit from the local tract alignment information.

### 2.5 Implementation and Parameter Settings

Our method is implemented using Pytorch (v1.9) [62] based on the existing VoxelMorph implementation^1^. Adam [63] is used for optimization with a learning rate of 10-4 as suggested in VoxelMorph [54]. For the overall network and each tract-specific backbone subnetwork, we train the model using a total of 50,000 batches (sufficient for achieving training and validation loss convergence), where each batch consists of one pair of input 3D volumes, i.e., a pair of 3D FA (1 channel per voxel) volumes or a pair of 3D tract-specific TOM (3 channels per voxel) volumes. For the weighting parameters, we set *λ_FA_* = 1, *λ_TOM_* = 1, and *λ_smooth_* = 0.01, which are the suggested settings for image registration (using MSE) and smoothing weighting parameters in the VoxelMorph package. Our unpublished parameter tuning experiments on a subset of the datasets (10 datasets) confirm that the selected parameter settings give in general the best registration performance. Our code will be open-source and the trained CNN registration model will be made available online^2^, as part of the SlicerDMRI project [64], [65].

## 3 Experimental Datasets

We demonstrate our method using dMRI datasets from multiple independently acquired populations, including: *1*) 100 young healthy adults (age: 29.1 ± 3.7 years; gender: 54 females and 46 males) in the Human Connectome Project (HCP) database [66], *2*) 30 teenagers (age: 10.2 ± 0.6 years; gender: 16 females and 14 males) in the Adolescent Brain Cognitive Development (ABCD) database [67], and *3*) 30 elderly adults (age: 63.1 ± 5.6 years; gender: 9 females and 21 males; 15 Parkinson’s disease patients and 15 healthy individuals) in the Parkinson’s Progression Markers Initiative (PPMI) database [68]. These dMRI datasets were scanned with different diffusion imaging protocols (see below Section 3.1 for acquisition details). In our study, the high-quality HCP datasets are used to train the registration model (n=50), as well as for model validation (n=20) and testing (n=30). The ABCD and PPMI datasets, where dMRI data were obtained with a clinical acquisition protocol from studies geared towards clinical applications, are used to test how the trained model generalizes to data from different populations and acquired using different acquisition protocols and scanners.

### 3.1 Data acquisition protocols

The HCP data were acquired with a high quality image acquisition protocol using a customized Connectome Siemens Skyra scanner [66]. The acquisition parameters are TE = 89.5 ms, TR = 5520 ms, phase partial Fourier = 6/8, voxel size = 1.25 × 1.25 × 1.25 mm^3^, and matrix = 145 × 174 × 145. A total of 288 images were acquired for each subject, including 18 baseline images and 270 diffusion weighted images evenly distributed at three shells of b = 1000/2000/3000 s/mm^3^. The dMRI data provided by HCP was processed following the well-designed HCP minimum processing pipeline [66], including brain masking, motion correction, eddy current correction, EPI distortion correction, and rigid registration to the MNI space.

The ABCD data included 30 dMRI datasets that were randomly selected, 10 per site, from three ABCD project scanner sites [67]. 20 subjects were scanned using a Siemens Prisma 3T Scanner, and 10 were scanned using a GE MR750 3T scanner. The dMRI data acquisition parameters are TE = 88 ms (Siemens) / 81.9 ms (GE), TR = 4100 ms (both), and voxel size = 1.7 × 1.7 × 1.7 mm^3^ (both). A total of 97 volumes were used for each dataset, including 7 baseline volumes with b = 0 s/mm^2^, 15 volumes with b = 1000 s/mm^2^, 15 volumes with b = 2000 s/mm^2^, and 60 volumes with b = 3000 s/mm^2^. The dMRI data provided by ABCD was processed as described in [69], including eddy current distortion correction, motion correction and B0 distortion correction.

The PPMI data were acquired using a 3T Siemens Trio scanner [68]. The dMRI data acquisition parameters are TE = 88 ms, TR = 7600 ms, and voxel size = 2 × 2 × 2 mm^3^. A total of 65 volumes were acquired for each subject, including 1 baseline image with b= 0 s/mm^2^ and 64 volumes with b = 1000 s/mm^2^. The dMRI data was processed as described in our previous study [70] using a well-established pipeline^3^, including eddy current-induced distortion correction, motion correction, and echo-planar imaging EPI distortion correction.

### 3.2 Data preprocessing

We performed the following preprocessing steps. First, each dMRI dataset from the ABCD and PPMI data was rigidly registered into the MNI space, i.e., where the HCP data for model training are, and upsampled to 1.25mm^3^ resolution (the same resolution as the HCP data). The rigid registration and upsampling are performed following the suggestion in the TractSeg software package^4^ for better computation of the tract segmentation and TOMs (which are used as method input and/or for evaluation, as described in the below paragraph). Then, each dMRI dataset in the HCP, ABCD and PPMI is cropped to 128 × 160 × 128 voxels to fit to our network input size, while ensuring coverage of the entire brain.

From each dMRI dataset, we compute the following data that is needed as input to our proposed method as well as other comparison methods (see Sec. 4). We first compute diffusion tensor images using a least squares estimation, followed by computation of the FA image from the tensor image, using the SlicerDMRI extension^5^ [64], [65] in 3D Slicer^6^. We also calculate multi-shell multi-tissue constrained spherical deconvolution (CSD) data, followed by FOD peak extraction with a maximum number of three peaks per voxel, using MRtrix^7^ [71], [72]. The extracted FOD peaks are subsequently provided to TractSeg to compute TOMs of 72 white matter tracts [55]. In our method, to reduce the number of TOMs while keeping all fiber orientation information, we combine bilateral tracts (e.g., the left and right arcuate fasciculus tracts) into one TOM. In total, this produced 40 TOMs (see Supplementary Table S1 for the list of TOMs).

For evaluation purposes (see Sec. 4.1 for details of the evaluation metrics), we compute the binary segmentation mask of each white matter tract [73]. We also perform tractography on the TOMs to compute fiber streamlines for each tract [55]. The masks and streamline tractography are computed using TractSeg with the default parameters suggested by the software. In addition, we also compute a tissue segmentation of the entire brain into WM, GM and CSF from the dMRI data using a deep learning method [74].

## 4 Experimental Comparisons and Evaluation Metrics

We perform two experimental evaluations. First, we assess the effectiveness of the proposed dMRI registration framework. Second, we compare the proposed method to four state-of-the-art dMRI registration methods. In the rest of the section, we introduce evaluation metrics used in these experiments (Sec. 4.1), followed by the details of the experimental comparisons (Secs. 4.2 and 4.3).

### 4.1 Evaluation Metrics

We first evaluate the dMRI registration methods by assessing their performance on aligning anatomically corresponding brain structures. To this end, we quantify the volumetric overlap of segmented brain structures between moved and target subjects (currently one of the most widely used evaluation metrics in medical image registration tasks [54], [57]). The volumetric overlap is computed using the well-known Dice score [54], [57], [75], which ranges from 0 to 1; a higher value means better overlap. First, we evaluate the volume overlap of each white matter tract by computing the Dice overlap score of the binary tract segmentation masks between the moved and target subjects. Next, we evaluate the volume overlap of the tissue segmentations (i.e., GM, WM, and CSF) of the entire brain. The Dice overlap score is computed for each tissue type to evaluate the performance. In the rest of the paper, we refer to these two metrics as *tract Dice score* and *tissue Dice score.*

Finally, in order to assess fiber spatial alignment, we include an evaluation using the geometric distances between fiber streamlines after registration. We use a metric based on pairwise fiber streamline distances, as proposed in several dMRI registration studies [11], [15], defined as follows:

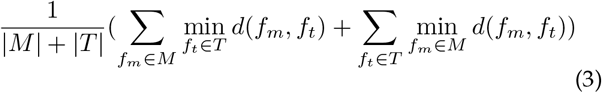

where *M* and *T* are the moved and target fiber tracts, *d* is the mean closest point fiber distance [7], [76] between two fibers in *M* and *T*, min_*f*_*i*∈*T*__ *d*(*f_m_, f_t_*) is the distance between the fiber *f_m_* and the fiber in *T* that is closest to *f_m_,* and, similarly, min_*f*_*i*∈*T*__ *d*(*f_m_, f_t_*) is the distance between the fiber *f_t_* and the fiber in *M* that is closest to *f_t_.* In the rest of the paper, we refer to this metric as *tract distance*. A lower tract distance represents a better spatial tract alignment.

### 4.2 Evaluation of the DDMReg Architecture

We first perform several ablation studies [77] to evaluate the effects of inclusion or exclusion of the proposed tract-specific backbone registration subnetworks. We compare our method with two baseline methods that do not use any tract-specific subnetworks. The first method registers only the FA image (referred to as Reg _FA_). Reg _FA_ uses the same U-net architecture as our whole-brain registration subnetwork, with a loss function that optimizes the MSE of FA and the smoothness of the deformation field (i.e., *L_FA_*(*f, m, ϕ*) = *λ_FA_L_FA_* + λ_*smooth*_ (*ϕ_wb_*)). The second baseline method uses both FA and TOM images (referred to as *Reg_FA+TOM_*). The same U-net architecture as in Reg_*FA*_ is used, but with a different loss function that includes an additional component to optimize the MSE of TOMs (i.e., 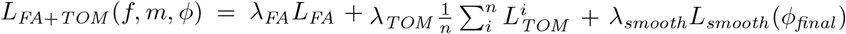). We use the same number of training batches and the same settings of the weighting parameters as our method (see Sec. 2.5).

We then assess the performance when including different numbers of tract-specific backbone registration subnetworks. In this experiment, we compare DDMReg (40 backbone subnetworks) with three additional methods that include only one randomly selected backbone subnetwork, three randomly selected backbone subnetworks, and twenty randomly selected backbone subnetworks (half of the total number of backbone networks). Here we also compare results to the Reg_*FA*_ method, which serves as a reference baseline method without any tract-specific backbone subnetworks.

Evaluation of the above methods is performed on the 30 HCP testing subjects. For each of the methods, between every pair of subjects (where 30 choose 2 gives a total of 435 pairs) we first perform registration and we then compute the evaluation metrics (tract Dice, tissue Dice and tract distance scores). Then, for each evaluation metric, we report the mean and the standard deviation across all subject pairs. A one-way repeated measures analysis of variance (ANOVA) is performed across the three methods for each evaluation metric, followed by a post-hoc paired t-test and effect size analysis (using Cohen’s *d* [78]) between each pair of the methods. In the assessment of different numbers of tract-specific backbones, we also separately report the mean tract Dice score and the mean tract distance for the tracts that are included or not included.

### 4.3 Comparison to State-of-the-Art Methods

We compare our proposed DDMReg method to four state-of-the-art dMRI registration methods, which are referred to as *SyN* [58] (which uses scalar FA images as input), *DTI-TK* [11] (which uses diffusion tensor images as input), *MRRegister* [34] (which uses FOD images as input), and *VoxelMorph* [54] (which uses scalar FA images and tract segmentations as input) methods in the rest of the paper. The experimental comparison is performed using the testing data including the 30 HCP, 30 ABCD and 30 PPMI subjects, which were acquired from different populations using different imaging protocols and scanners (see Section 3.1). For parameters involved in each of these compared methods, we use the default values as suggested in their software. All registration commands were run in the default way without specifying any parameter modifications.

The SyN method performs registration by optimizing a cross correlation similarity measure between scalar images [58] and it has been shown to be successful in several comparative studies [15], [79]. In particular, for dMRI registration, applying SyN on FA images has shown good registration performance (second-ranked across six compared methods [15].) In our study, we use the SyN implementation in Advanced Normalization Tools (ANTs) [80].

The DTK-TK method performs dMRI registration based on the diffusion tensor images, with the aim of optimizing the L2 inner product distance between the tensors [11]. In two comparative studies for dMRI registration [15], [81], DTK-TK achieved the best performance (including the aforementioned study where the SyN method ranked second [15].) In our study, we use the DTK-TK implementation in the Diffusion Tensor Imaging ToolKit^8^.

The MRRegister method is a more recently proposed FOD-based method that performs dMRI registration [34]. FOD-based methods aim to improve registration in regions of crossing fibers [32]–[35]. MRRegister optimizes a diffeomorphic non-linear transformation model based on the mean squared difference and cross-correlation of the FOD spherical harmonic coefficients. In our study, we use the implementation in MRtrix [36].

The VoxelMorph method is a deep learning method that learns a deformation field between pairs of images [54]. VoxelMorph is not designed specifically for dMRI registration and there has been no previous evaluation on dMRI data. Therefore, we test different combinations of inputs (see Supplementary Fig. S1) and choose the best performing input (i.e., FA) for use in this experiment. The VoxelMorph model is trained to register FA images by directly using the VoxelMorph package with default parameters. In addition, it is suggested in [54] that auxiliary information in the form of anatomical segmentations can be used for VoxelMorph registration improvement. (This is done by optimizing the Dice score of the segmentations [54].) For a complete VoxelMorph comparison, we therefore use the option in the VoxelMorph package to include an additional auxiliary loss for optimizing the Dice overlap score of the segmentations of the 40 white matter tracts. The implementation in the VoxelMorph package is used.

After obtaining the deformation fields, the binary tract segmentation masks and the TOMs of each moving subject are warped to the corresponding target subject space for computing the evaluation metrics. For reorientation of the TOMs, we use the vecreg command provided in FSL [82] (the recommended software for TOM reorientation suggested by the TractSeg package).

For experimental comparison, we perform registration between every pair of subjects across all testing data (30 HCP, 30 ABCD and 30 PPMI subjects), which we refer to the “all-to-all” evaluation in the rest of the paper. We report the mean and the standard deviation across all testing subject pairs for each evaluation metric. ANOVA, followed by a paired t-test and effect size analysis (using Cohen’s *d*) between each pair of compared methods, are used for statistical comparison. To access performance of each method on registering data from different populations and acquired with different protocols and scanners, we also report evaluation metrics of the intra-population subject pairs (i.e., between HCP and HCP, ABCD and ABCD, and PPMI and PPMI) and inter-population subject pairs (i.e., between HCP and ABCD, between HCP and PPMI, and between ABCD and PPMI). In addition, to evaluate the regularity of the deformation fields, we compute the Jacobian determinant of the deformation field of each testing image pair (as used in multiple related studies [54], [83], [84]). Specifically, the Jacobian matrix at each voxel captures local properties around the voxel, where a non-positive determinant indicates that the deformation is not diffeomorphic [54], [85]. In our study, we report the mean percentage of voxels with a non-positive Jacobian determinant [54] across all testing image pairs for each method. We also compute the bending energy [86] that has been widely used for accessing the smoothness of the deformation field in medical image registration tasks [87], [88]. In our study, the mean of the bending energy over the entire deformation (computed using implementation of bending energy loss in the MONAI project^9^) across all testing image pairs for each method is reported for each method. For each compared method, we also report the computational time for data preprocessing to prepare the method input and that for registering the image pair to estimate the deformation field.

We also perform an evaluation to estimate registration uncertainty given the deformation fields from the registration subnetworks. Uncertainty estimates can provide intuitive information about the quality of image registration [89]. In our study, given the fact that multiple tracts can cross at the same voxel, uncertainty estimates are important to show if there are tract-specific effects. To quantify registration uncertainty, for each voxel we compute the covariance matrix of the estimated displacement vectors (a 3×3 matrix) across the 40 tract-specific deformation fields. For visualization of the uncertainty, we calculate the mean of the square root of the diagonal values of the covariance matrix for each voxel. A lower value indicates higher agreement across the registration fields and thus high registration certainty.

## 5 Experimental Results

### 5.1 Evaluation of the DDMReg Architecture

As shown in Fig. 3, using only FA for registration (i.e., Reg_*FA*_) generated the lowest mean tract Dice score, the lowest mean tissue Dice score, and the highest tract distance (0.734, 0.710, and 3.724 mm, respectively) after registration. Adding an additional loss component to penalize differences of the TOMs (i.e., Reg_*FA+TOM*_) improved these evaluation metrics (0.766, 0.734, and 3.573 mm, respectively). Further including the tract-specific backbone registration subnetworks (*Reg_Proposed_*) achieved the best results (0.786, 0.739, and 3.418mm, respectively). For each of the evaluation metrics, the ANOVA analysis showed a statistically significant difference in the three-method comparison (*p* < 0.001), where *Reg_Proposed_* had significantly higher tract Dice, higher tissue Dice, and lower tract distance scores than the other two methods (*p* < 0.001 and Cohen’s *d* > 1). See Supplementary Table S2 for detailed statistical comparison results. The detailed evaluation metrics for each tract and each tissue are provided in Supplementary Fig. S2.

**Fig. 3.**
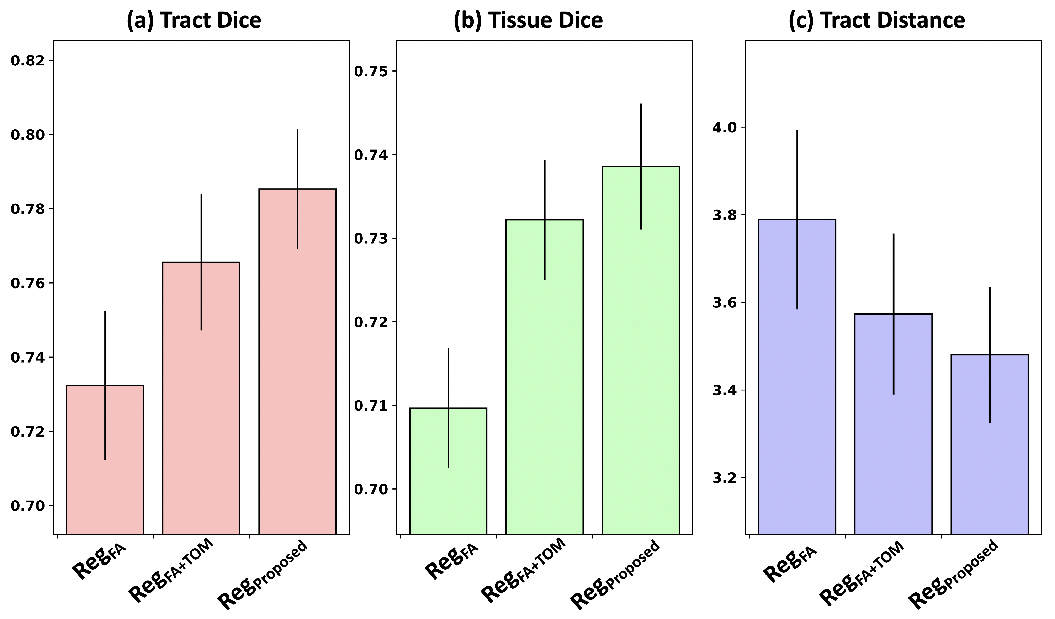
Comparison between different network inputs and loss functions. Compared to the baseline methods (without any tract-specific subnetworks), the results of this ablation study show that the proposed DDM-Reg architecture using pretrained tract-specific backbone registration subnetworks improves the registration performance.

Fig. 4 shows the comparison using different numbers of tract-specific backbone registration subnetworks. From subfigures 4(a) to 4(c), we can observe that including more backbones increased the registration performance in particular in the tract-wise evaluation metrics (tract Dice and tract distance). However, the whole brain evaluation metric (tissue Dice) increased only slightly after using more than three backbones. This is because tissue Dice evaluates the performance of large anatomical regions (the entire WM, GM and CSF) and does not consider local white matter structure alignment of individual tracts. Compared to the method that does not use any backbones, including all backbones (n = 40) increases the tract Dice score by 0.052, the tissue Dice score by 0.029, and decreases the tract distance by 0.306 mm. In addition, from subfigures 4(d) and 4(e), we can observe that including more backbone tracts increased the performance on registering the tracts that were not included during training.

**Fig. 4.**
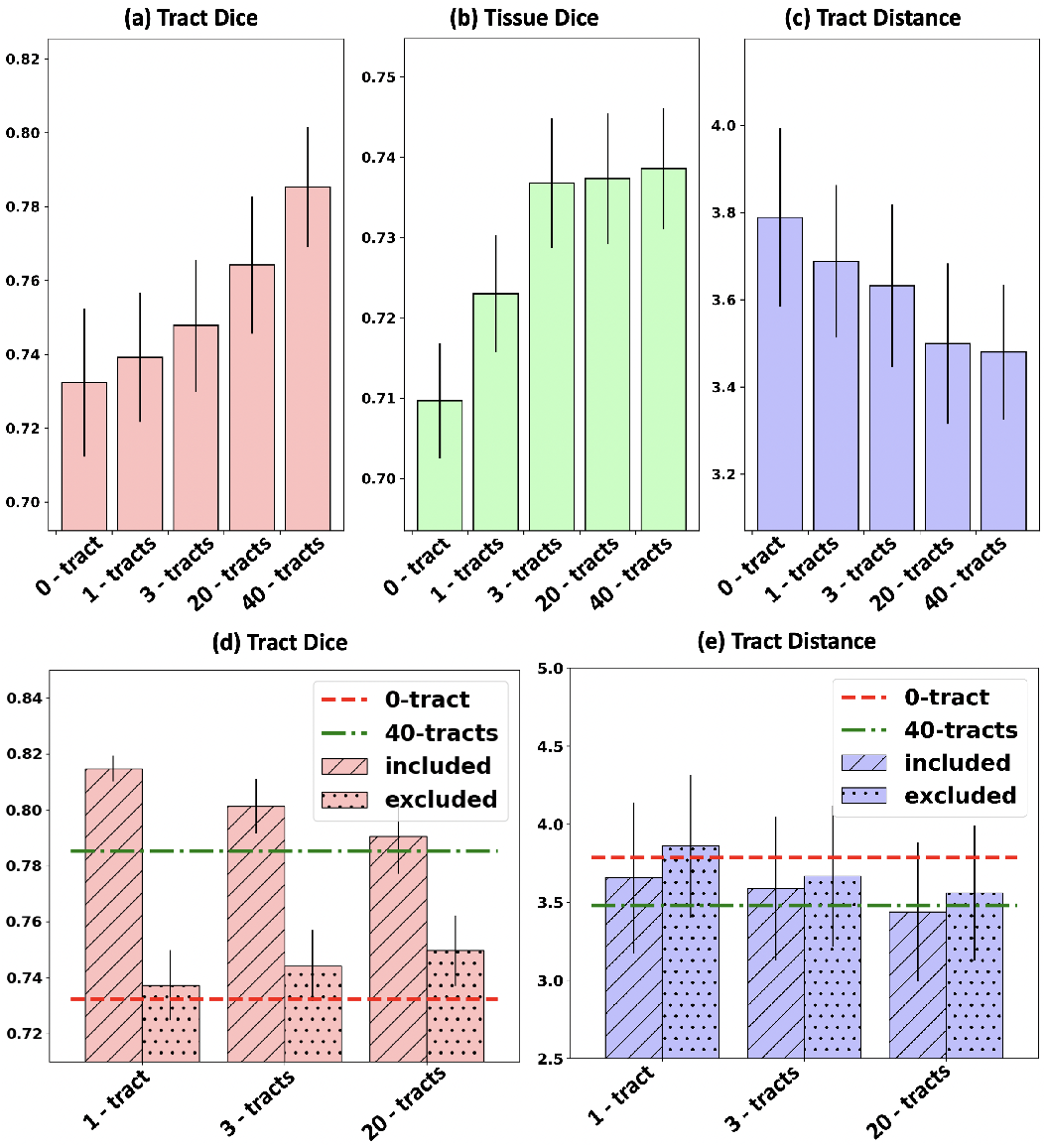
Comparison across different numbers of backbone tracts. In general, the registration performance is improved when including more backbone tracts.

### 5.2 Comparison to State-of-the-Art Methods

As shown in Fig. 5, the proposed DDMReg method obtained overall the best results (the highest tract Dice, the highest tissue Dice, and the lowest tract distance) across all the compared methods in the “all-to-all” evaluation. DTI-TK, VoxelMorph and SyN were the second ranking methods for tract Dice, tissue Dice and tract distance, respectively, while MRRegister had the least favored performance except for the tissue Dice where MRRegister obtained a slightly higher score than DTI-TK. For each of the evaluation metrics, the ANOVA analysis showed a significant difference in the five-method comparison (*p* < 0.001), where DDMReg had significantly higher tract Dice, higher tissue Dice, and lower tract distance scores than the other four compared methods (*p* < 0.001 and Cohen’s *d* ≥ 0.9). See Supplementary Table S3 for detailed statistical comparison results. The detailed evaluation metrics for each tract and each tissue are provided in Supplementary Fig. S3. In the evaluation of the registration of intra- and inter-population subject pairs, DDMReg also achieved the best results in each of the six categories (i.e., within HCP, within ABCD, within PPMI, between HCP and ABCD, between HCP and PPMI, and between ABCD and PPMI), except in registration between HCP and PPMI where DTI-TK obtained a slightly higher tract Dice than DDMReg (0.7475 versus 0.7469). The performance of the other compared methods varied across the datasets and the evaluation metrics.

**Fig. 5.**
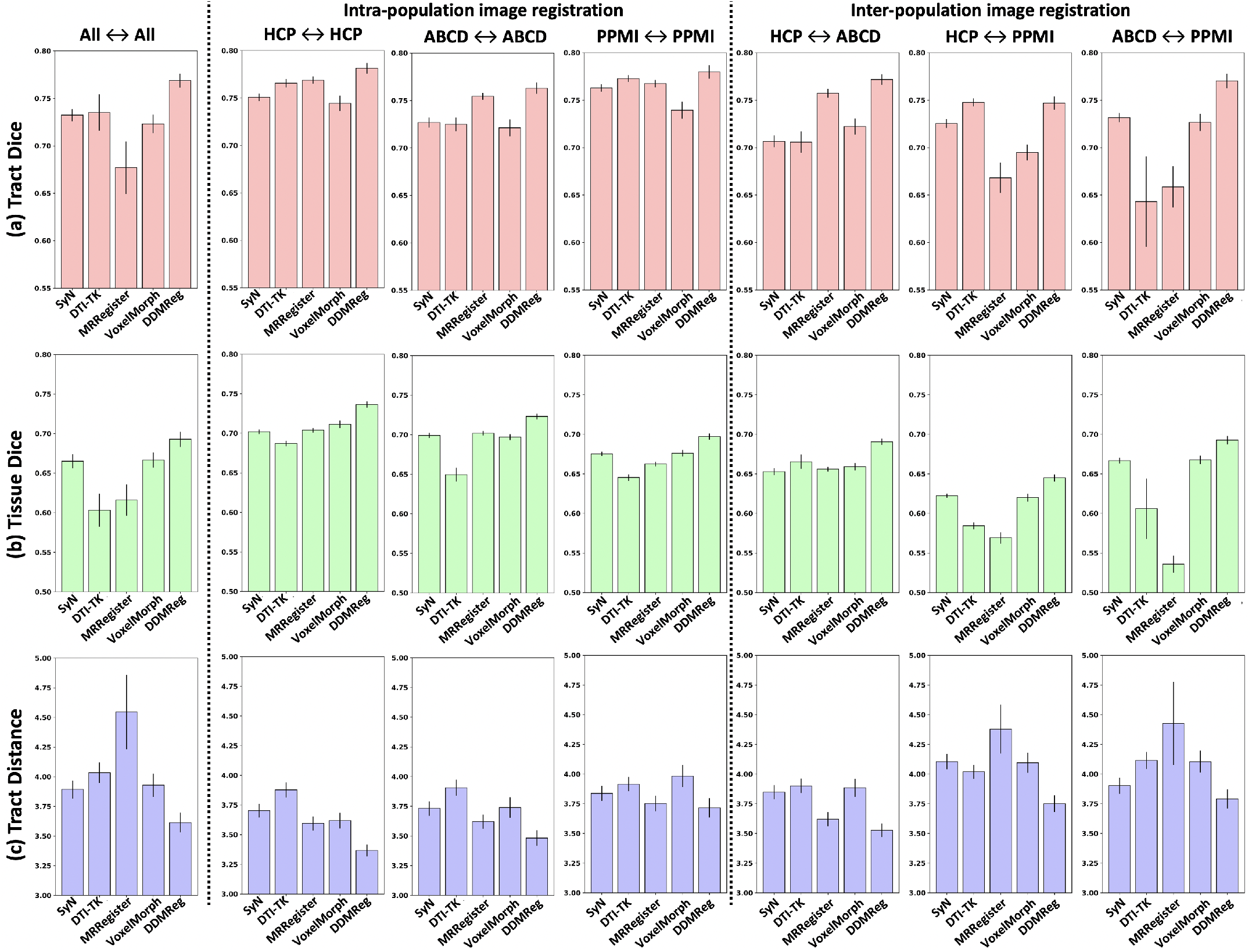
Comparison to the state-of-the-art methods. The first column shows the comparison results from registering image pairs across all testing datasets (i.e., the “all-to-all” evaluation), the second to the fourth columns show the results from registering image pairs from the same population and scanned using the same imaging protocol, and the final three columns show the results from registering image pairs from different populations and different scanners. Overall, the proposed DDMReg method achieves the best performance. DDMReg obtains the highest tract Dice score, the highest tissue Dice score, and the lowest tract distance in the “all-to-all” evaluation as well as in the intra- and inter-population evaluations, except in registration between HCP and PPMI where DTI-TK obtained a slightly higher tract Dice than DDMReg (0.7475 versus 0.7469).

Fig. 6 gives a visualization of registration results for each of the compared methods. Two example image pairs are selected, including: (1) one randomly selected HCP subject pair, for a demonstration of registering datasets scanned using the same imaging protocol and from the same population, and (2) one subject pair of the youngest teenager in the ABDC testing data (9 years old) and the oldest elderly adult in the PPMI testing data (72 years old), for a demonstration of registering datasets from different populations and scanners. In example (1), all methods generate visually good results to align the two brains, while we can observe differences in local regions. As shown in the inset images that render the anterior part of the anterior thalamic radiation (ATR) tract, VoxelMorph and DDMReg obtain visually more similar results to the target image than the SyN, DTI-TK and MRRegister methods. In example (2), SyN and DDMReg generate a visually better alignment of ABCD and PPMI data compared to the DTI-TK, MRRegister and VoxelMorph methods, in particular in the local white matter region near the fronto-pontine tract (FPT). Compared to the SyN method, DDMReg obtains a visually better FPT segmentation that is more similar to the target PFT segmentation. See Supplementary Fig. S4 for additional visualization of the above two examples, and Supplementary Fig. S5 for visualization of more example image pairs from different datasets.

**Fig. 6.**
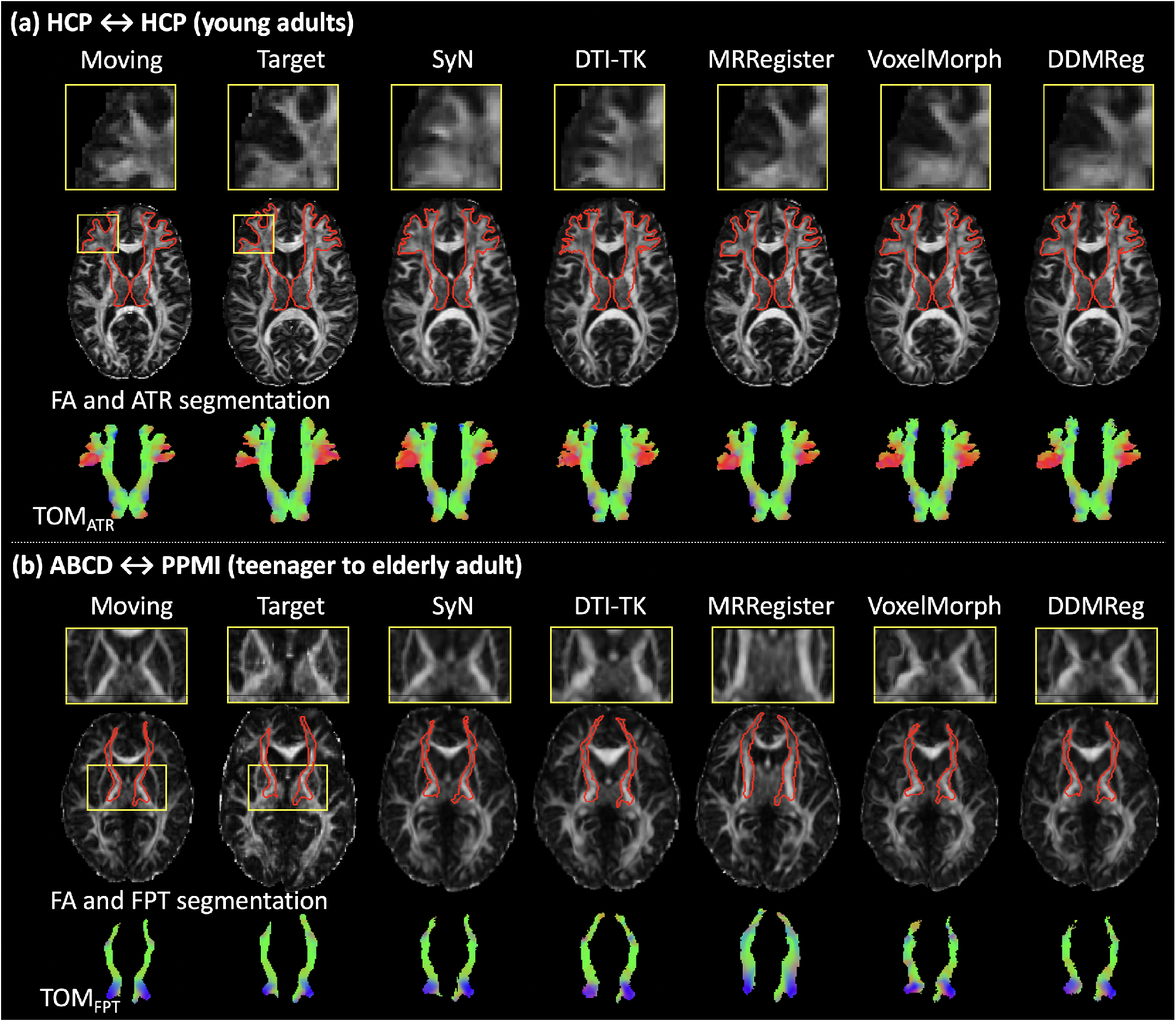
Visualization of registration results for the state-of-the-art and DDMReg methods. Two selected example image pairs are shown. Wholebrain FA images are displayed, plus insert images giving a zoomed-in visualization for a part of the Anterior Thalamic Radiation (ATR) in subfigure (a) and the fronto-pontine tract (FPT) in subfigure (b). The segmentation of the entire ATR and FPT (overlaid on the FA image in red) and the TOM (shown in directionally encoded color [90]) are also provided.

Fig. 7 gives a visualization result of the proposed DDM-Reg method, including the FA images, the estimated deformation field, and the registration uncertainty map. The same example image pairs as used in Fig. 6 are used. We can observe a good spatial alignment of the FA images after registration. For example, the sizes of the brains (more apparent in the axial view), the shapes of the lateral ventricle (red arrows in the coronal view), and the shapes of the corpus callosum (green arrows in the sagittal view) are highly visually similar between the target and the moved images. The deformation field is visually smooth, showing a regular, smooth displacement after registration. (We refer the readers to Fig. 10 in [54] for a visualization of regular and irregular deformation fields.) The deformation field and uncertainty map show no tract-specific effects (e.g., no particular tract regions showing high uncertainty), even though our tract-specific registration subnetworks register based on individual tracts only. In the example HCP subject pair, the uncertainty is generally highest near the cortex, where anatomical variability across subjects is greater, and lowest in the deep white matter. In the example PPMI and ABCD image pair, the uncertainty is relatively higher than that in the HCP data example, because it is a more difficult task to register teenage and elderly adult brains that were acquired using different protocols and scanners. See Supplementary Figs. S6 and S7 for additional visualizations of registration results of the proposed method.

**Fig. 7.**
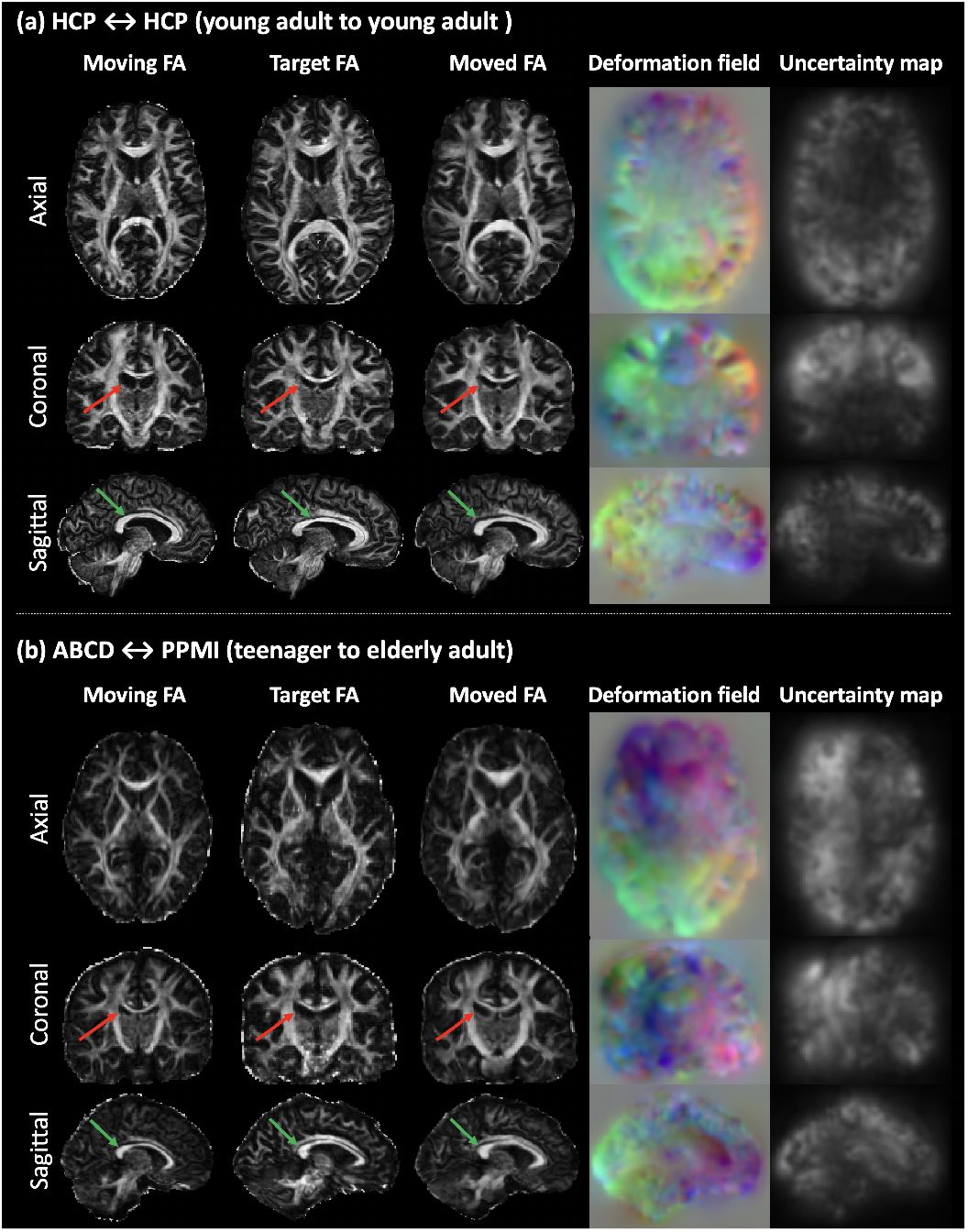
Visualization of registration results of the proposed DDMReg method, including the FA images, the deformation field and the registration uncertainty map. The red and green arrows indicate the lateral ventricle and the corpus callosum.

#### Jacobian determinant and bending energy

For all methods, the percentage of voxels with a non-positive Jacobian determinant is very low (fewer than 0.01%) except for SyN that is relatively high 0.05%, which is still a very low percentage. The bending energies of the SyN, DTI-TK, MRRegister, VoxelMorph and DDMReg methods are 0.010, 0.051, 0.067, 0.023, and 0.018, respectively, where all methods have relatively similar and low values. These indicate that all methods produce good deformation regularity and smoothness.

#### Runtime

Table 1 gives the computational time for registering a pair of example HCP dMRI datasets for each method. The computation is performed on a Linux workstation, equipped with 8 DIMMs for CPU computation (8×32GB memory) and 4 NVIDIA GeForce GTX 1080 Ti Graphics Cards for GPU computation (4× 11GB memory). For each method, if multi-thread CPU computing is supported, we use 16 threads. The inputs to SyN, DTI-TK and VoxelMorph methods are based on DTI; thus these methods take a small amount of time for computing the input data. The MRRegister and DDMReg methods require the computation of CSD data and FOD peaks, which take a relatively long time. For registering the input data, the SyN, DTI-TK, and MRRegister methods are based on the traditional image registration strategy and can only use CPU computation; thus, they require a relatively long time to register. The VoxelMorph and our proposed DDMReg methods can leverage the GPU, thus highly reducing the computational time. We note that the computational time of VoxelMorph and DDMReg is also small when only using the CPU.

**TABLE 1.** Computational time (in seconds).

| Method | CPU/GPU | Preprocess | Register |
| --- | --- | --- | --- |
| SyN | CPU | 60s (FA) | 350s |
| DTI-TK | CPU | 60s (Tensor) | 950s |
| MRRegister | CPU | 1400s (FOD) | 350s |
| VoxelMorph | CPU | 60s (FA) | 10s |
|  | CPU+GPU |  | 5s |
| DDMReg | CPU | 1800s (FA+FOD+TOM) | 80s |
|  | CPU+GPU | 1600s (FA+FOD+TOM) | 10s |

## 6 Discussion

In this work, we have proposed a novel deep learning method, DDMReg, that uses joint whole-brain and tract-specific information for dMRI registration. DDMReg is an unsupervised method for deformable registration between pairs of dMRI datasets. We proposed a novel registration architecture that leverages not only whole brain information but also tract-specific fiber orientation information. We compared DDMReg to several state-of-the-art methods, and we showed highly improved registration performance. Below, we discuss several detailed observations about the experimental results.

We designed a novel network architecture for dMRI registration, which enables the ensemble of multiple registration subnetworks trained on different types of input data to form one unified prediction. This is similar to the concept of ensemble learning which has been shown to be highly successful in many deep learning tasks [91]–[95]. In our method, we leveraged the successful U-net-based registration architecture proposed in VoxelMorph [54], which was found to be highly effective for registration of individual types of input data (either FA or TOM). VoxelMorph has also been shown to be successful for registration tasks that include other types of dMRI data as input such as diffusion tensor images [53]. Then, based on our recently proposed dual-stream network that demonstrated success to combine two deformation fields for multimodal image registration [57], our proposed multi-deformation fusion subnetwork combined the outputs of the multiple U-net registration subnetworks. This enabled training the overall network in such a way that the registration subnetworks constrained each other, while cooperating to optimize for a unified registration deformation field. In our experiments, we demonstrated that the ensemble of the whole-brain and the tract-specific registration subnetworks achieved reciprocal advantages to both types of the input data, i.e., improved performance on both whole-brain-wise (tissue Dice score) and tract-specific (tract Dice score and tract distance) evaluation metrics.

We demonstrated advantages by using pretrained backbone registration networks that enabled the usage of multiple registration subnetworks together. In the literature, backbone networks are often trained for a specific task and later used by fine tuning to another networks that may solve a different task. Pretrained backbones have been widely used in deep learning for medical image computing tasks and have been shown to improve both effectiveness and efficiency [59]–[61]. In our work, we pretrained registration networks to perform tract-specific registration, which are later used as backbones for registration of the entire dMRI data. Leveraging pretrained backbones can reduce the computational burden, so that we were able to use a large number of U-nets (41 in our study) in the same network. Our results demonstrated that as the number of included pretrained backbones increased, the registration performance kept increasing. We highlight that our proposed pretrained backbone strategy can also enable inclusion of additional inputs, such as T1w, T2w and functional MRI, for registration using multimodal imaging information.

Our method benefited from the unique tract-specific fiber orientation information encoded in the TOM. TOMs have been shown to be successful for tract-specific tractography [55], demonstrating their ability to encode fiber geometric information. Our results showed that including TOMs for dMRI registration greatly improved spatial tract alignment between different subjects after registration. In addition, each voxel in a TOM has only one FOD peak specific to the fiber orientation from a certain white matter tract. Due to this, TOMs are able to unambiguously model crossing fibers (where multiple white matter tracts with different fiber orientations are present within the same voxel). This has two main advantages. First, using TOMs can improve registration in the crossing fiber regions that are widespread in the white matter anatomy. Second, using TOMs can solve the problem that the FOD peak correspondence across subjects is not known when multiple FOD peaks are present in one voxel. However, it should be mentioned that TOMs are derived from tractography, which is not a perfect representation of the true underlying fiber tract anatomy. The CNN that produces TOMs was trained using fiber orientations estimated from tractography streamlines [55]. While such streamlines are highly consistent across subjects [70] and are useful for registration [41]–[45], tractography can suffer from false positive and false negative fiber errors [40], [96]. If such errors affected training of the TOMs, biases could potentially have been introduced. We believe the effect of any biases on our registration method should be small, for three reasons. First, the training of TOMs leveraged a well-curated dataset designed to reduce potential tractography errors by employing fiber filtering and expert refinement [55], [73]. Second, the training dataset included a high number of streamlines per tract from multiple subjects, reducing the effect of errors in individual streamlines. Third, in addition to TOMs, our registration method benefits from FA microstructure images, which are computed directly from the dMRI data and do not rely on tractography.

We showed a good generalizability of the proposed DDMReg to dMRI data from different populations with varying ages and acquired using different imaging protocols and different scanners. (The ability of a dMRI registration method to generalize to data from different acquisitions is important, because dMRI acquisitions can have widely varying numbers of gradient directions, b-values, and magnitude of b-values, posing a challenge for machine learning.) Several aspects of our method contribute to the good generalizability. In training, our method benefits from high-quality multi-shell HCP data. The strategy of training on HCP data for application to other datasets has been successful in multiple dMRI machine learning approaches for tasks such as brain tissue segmentation [74] and white matter tract parcellation [73]. In addition, our method benefits from the good generalizability of TOMs that are well-suited to represent dMRI data acquired with different scanners, different spatial and angular resolutions, and different b-values [55]. Finally, our method incorporates the FA image, which is widely used and successful for dMRI registration [5], [15]–[20].

The proposed DDMReg method demonstrated advanced dMRI registration performance compared to several state-of-the-art methods. DDMReg leverages the FA image, which provides information about the tissue microstructure of the brain as a whole, and it incorporates TOMs, which are effective to describe fiber orientation information, in particular in the presence of crossing fibers. Two of the compared methods, SyN and VoxelMorph, obtained generally good performance in the overall “all-to-all” evaluation by performing registration based on FA images. The other two compared methods, DTI-TK and MRRegister, performed generally well on the intra-population registration experiments, but their performance suffered in the interpopulation experiments, especially when registering data between ABCD and PPMI datasets (teenagers and elderly adults). DTI-TK and MRRegister used diffusion tensor data and FOD data as input, respectively. These input data types can capture fiber orientation information, but may be more challenging to register when considering data from populations with different ages and/or scanned with different imaging protocols and scanners.

Potential limitations of the present study, including suggested future work to address limitations, are as follows. First, our proposed method performs a pair-wise registration between dMRI datasets. An interesting future investigation includes extending our method to enable groupwise registration for anatomical template creation. Second, in the present study, we used all TOMs that are available in TractSeg. We demonstrated that as the number of included TOMs increased, the registration performance kept increasing. Therefore, it would be interesting to include additional TOMs that are not currently available in TractSeg. Third, currently the computation of a TOM is relatively time-consuming because of the computation of CSD and FOD peaks. A future investigation could include deep learning to perform CSD computation to further improve computational speed [97]. Fourth, while we demonstrated a high generalization of DDMReg to dMRI data from different populations including teenagers, young and elderly healthy adults and Parkinson’s disease patients, future work could include an investigation of brains with more challenging pathologies, e.g., brain tumors. Such work could require curation of training and testing data from patients, as well as possible changes to the network architecture (e.g., to consider lesion segmentations) and/or algorithm parameters (e.g., loss function weighting parameters).

## 7 Conclusion

In this paper, we have presented DDMReg, a novel deep learning method that leverages joint whole-brain and tract-specific information for accurate dMRI registration. Experimental results show that DDMReg achieves significantly improved registration accuracy compared to several state-of-the-art methods. Importantly, we demonstrate successful generalization of DDMReg to dMRI data from different populations with varying ages and acquired using different acquisition protocols and different scanners.

## Acknowledgment

We would like to thank Dr. Lipeng Ning and Jianzhong He for their technical support. We also gratefully thank the TractSeg group and the VoxelMorph group for making their code and/or trained deep learning models used in this study available online. We acknowledge funding provided by the following National Institutes of Health (NIH) grants: P41 EB015902, P41 EB015898, R01 MH074794, R01 MH125860, P41 EB028741, and R01 MH119222.

## Footnotes

1. github.com/voxelmorph/voxelmorph

2. github.com/SlicerDMRI/DDMReg

3. github.com/pnlbwh/pnlutil

4. github.com/MIC-DKFZ/TractSeg

5. dmri.slicer.org

6. www.slicer.org

7. www.mrtrix.org

8. http://dti-tk.sourceforge.net/pmwiki/pmwiki.php

9. github.com/Project-MONAI/MONAI/blob/dev/monai/losses/deform.py

## References

[1] P. Basser, J. Mattiello, and D. LeBihan, “MR diffusion tensor spectroscopy and imaging,” Biophys. J., vol. 66, no. 1, pp. 259–267, 1994.

[2] P. Basser, S. Pajevic, C. Pierpaoli, J. Duda, and A. Aldroubi, “In vivo fiber tractography using DT-MRI data,” Mag. Res. Med., vol. 44, no. 4, pp. 625–632, 2000.

[3] C. Pierpaoli and P. Basser, “Toward a quantitative assessment of diffusion anisotropy,” Mag. Res. Med., vol. 36, no. 6, pp. 893–906, 1996.

[4] S. Cetin-Karayumak, M. Di Biase, N. Iturry, B. Reid, N. Somes, A. Lyall et al., “White matter abnormalities across the lifespan of schizophrenia: A harmonized multi-site diffusion MRI study,” Mol. Psychiatry, vol. 25, p. 3208–3219, 2020.

[5] E. Moulton, R. Valabregue, B. Díaz, C. Kemlin, S. Leder, S. Lehéricy et al., “Comparison of spatial normalization strategies of diffusion MRI data for studying motor outcome in subacute-chronic and acute stroke,” NeuroImage, vol. 183, pp. 186–199, 2018.

[6] A. Varentsova, S. Zhang, and K. Arfanakis, “Development of a high angular resolution diffusion imaging human brain template,” NeuroImage, vol. 91, pp. 177–186, 2014.

[7] F. Zhang, Y. Wu, I. Norton, L. Rigolo, Y. Rathi, N. Makris et al., “An anatomically curated fiber clustering white matter atlas for consistent white matter tract parcellation across the lifespan,” NeuroImage, vol. 179, pp. 429–447, 2018.

[8] F.-C. Yeh, S. Panesar, D. Fernandes, A. Meola, M. Yoshino, J. C. Fernandez-Miranda et al., “Population-averaged atlas of the macroscale human structural connectome and its network topology,” NeuroImage, vol. 178, pp. 57–68, 2018.

[9] J. Du, A. Goh, and A. Qiu, “Diffeomorphic metric mapping of high angular resolution diffusion imaging based on riemannian structure of orientation distribution functions,” IEEE TMI, vol. 31, no. 5, pp. 1021–1033, 2011.

[10] D. C. Alexander, C. Pierpaoli, P. J. Basser, and J. C. Gee, “Spatial transformations of diffusion tensor magnetic resonance images,” IEEE TMI, vol. 20, no. 11, pp. 1131–1139, 2001.

[11] H. Zhang, P. Yushkevich, D. Alexander, and J. Gee, “Deformable registration of diffusion tensor MR images with explicit orientation optimization,” MedIA, vol. 10, no. 5, pp. 764–785, 2006.

[12] J. Duarte, G. Sapiro, N. Harel, and C. Lenglet, “A framework for linear and non-linear registration of diffusion-weighted MRIs using angular interpolation,” Front. Neuroscience, vol. 7, p. 41, 2013.

[13] P. Zhang, M. Niethammer, D. Shen, and P.-T. Yap, “Large deformation diffeomorphic registration of diffusion-weighted imaging data,” MedIA, vol. 18, no. 8, pp. 1290–1298, 2014.

[14] L. J. O’Donnell, A. Daducci, D. Wassermann, and C. Lenglet, “Advances in computational and statistical diffusion MRI,” NMR in Biomedicine, vol. 32, no. 4, p. e3805, 2019.

[15] Y. Wang, Y. Shen, D. Liu, G. Li, Z. Guo, Y. Fan et al., “Evaluations of diffusion tensor image registration based on fiber tractography,” Biomedical engineering online, vol. 16, no. 1, pp. 1–20, 2017.

[16] S. M. Smith, M. Jenkinson, H. Johansen-Berg, D. Rueckert, T. E. Nichols, C. E. Mackay et al., “Tract-based spatial statistics: voxelwise analysis of multi-subject diffusion data,” NeuroImage, vol. 31, no. 4, pp. 1487–1505, 2006.

[17] P.-H. Yeh, K. Simpson, T. C. Durazzo, S. Gazdzinski, and D. J. Meyerhoff, “Tract-Based Spatial Statistics (TBSS) of diffusion tensor imaging data in alcohol dependence: abnormalities of the motivational neurocircuitry,” Psychiatry Research: Neuroimaging, vol. 173, no. 1, pp. 22–30, 2009.

[18] M. Malinsky, R. Peter, E. Hodneland, A. J. Lundervold, A. Lundervold, and J. Jan, “Registration of FA and T1-weighted MRI data of healthy human brain based on template matching and normalized cross-correlation,” J. Dig. Imag., vol. 26, no. 4, pp. 774–785, 2013.

[19] F. Zhang, Y. Wu, I. Norton, Y. Rathi, A. J. Golby, and L. J. O’Donnell, “Test–retest reproducibility of white matter parcellation using diffusion MRI tractography fiber clustering,” Hum. Brain Mapp., vol. 40, pp. 3041–3057, 2019.

[20] B. Li, W. J. Niessen, S. Klein, M. de Groot, M. A. Ikram, M. W. Vernooij et al., “Longitudinal diffusion MRI analysis using Segis-Net: a single-step deep-learning framework for simultaneous segmentation and registration,” NeuroImage, vol. 235, p. 118004, 2021.

[21] D. C. Alexander and J. C. Gee, “Elastic matching of diffusion tensor images,” CVIU, vol. 77, no. 2, pp. 233–250, 2000.

[22] J. Ruiz-Alzola, C.-F. Westin, S. K. Warfield, C. Alberola, S. Maier, and R. Kikinis, “Nonrigid registration of 3D tensor medical data,” MedIA, vol. 6, no. 2, pp. 143–161, 2002.

[23] A. Leemans, J. Sijbers, S. DeBacker, E. Vandervliet, and P. Parizel, “Affine coregistration of diffusion tensor magnetic resonance images using mutual information,” in ACIVS, 2005, pp. 523–530.

[24] Y. Cao, M. I. Miller, R. L. Winslow, and L. Younes, “Large deformation diffeomorphic metric mapping of vector fields,” IEEE TMI, vol. 24, no. 9, pp. 1216–1230, 2005.

[25] B. Yeo, T. Vercauteren, P. Fillard, J. Peyrat, X. Pennec, P. Golland et al., “DT-REFinD: Diffusion tensor registration with exact finite-strain differential,” IEEE TMI, vol. 28, no. 12, pp. 1914–1928, 2009.

[26] P.-T. Yap, G. Wu, H. Zhu, W. Lin, and D. Shen, “TIMER: Tensor image morphing for elastic registration,” NeuroImage, vol. 47, no. 2, pp. 549–563, 2009.

[27] B. Jeurissen, A. Leemans, J.-D. Tournier, D. K. Jones, and J. Sijbers, “Investigating the prevalence of complex fiber configurations in white matter tissue with diffusion magnetic resonance imaging,” Hum. Brain Mapp., vol. 34, no. 11, pp. 2747–2766, 2013.

[28] A. Barmpoutis, B. C. Vemuri, and J. R. Forder, “Registration of high angular resolution diffusion MRI images using 4 th order tensors,” in MICCAI, 2007, pp. 908–915.

[29] M.-C. Chiang, A. D. Leow, A. D. Klunder, R. A. Dutton, M. Barysheva, S. E. Rose et al., “Fluid registration of diffusion tensor images using information theory,” IEEE TMI, vol. 27, no. 4, pp. 442–456, 2008.

[30] G. Cheng, B. Vemuri, P. Carney, and T. Mareci, “Non-rigid registration of high angular resolution diffusion images represented by gaussian mixture fields,” in MICCAI, 2009, pp. 190–197.

[31] X. Geng, T. J. Ross, W. Zhan, H. Gu, Y.-P. Chao, C.-P. Lin et al., “Diffusion MRI registration using orientation distribution functions,” in IPMI, 2009, pp. 626–637.

[32] L. Bloy and R. Verma, “Demons registration of high angular resolution diffusion images,” in ISBI, 2010, pp. 1013–1016.

[33] X. Hong, L. R. Arlinghaus, and A. W. Anderson, “Spatial normalization of the fiber orientation distribution based on high angular resolution diffusion imaging data,” Mag. Res. Med., vol. 61, no. 6, pp. 1520–1527, 2009.

[34] D. Raffelt, J. Tournier, J. Fripp, S. Crozier, A. Connelly, and O. Salvado, “Symmetric diffeomorphic registration of fibre orientation distributions,” NeuroImage, vol. 56, no. 3, pp. 1171–1180, 2011.

[35] Y. Qiao, W. Sun, and Y. Shi, “FOD-based registration for susceptibility distortion correction in brainstem connectome imaging,” NeuroImage, vol. 202, p. 116164, 2019.

[36] J.-D. Tournier, R. Smith, D. Raffelt, R. Tabbara, T. Dhollander, M. Pietsch et al., “MRtrix3: A fast, flexible and open software framework for medical image processing and visualisation,” NeuroImage, vol. 202, p. 116–137, 2019.

[37] E. Moulton, R. Valabregue, B. Diaz, C. Kemlin, S. Leder, S. Lehéricy et al., “Comparison of spatial normalization strategies of diffusion mri data for studying motor outcome in subacute-chronic and acute stroke,” NeuroImage, vol. 183, pp. 186–199, 2018.

[38] R. Dimitrova, M. Pietsch, D. Christiaens, J. Ciarrusta, T. Wolfers, D. Batalle et al., “Heterogeneity in brain microstructural development following preterm birth,” Cerebral Cortex, vol. 30, no. 9, pp. 4800–4810, 2020.

[39] K. Pannek, J. Fripp, J. M. George, S. Fiori, P. B. Colditz, R. N. Boyd et al., “Fixel-based analysis reveals alterations is brain microstructure and macrostructure of preterm-born infants at term equivalent age,” NeuroImage: Clinical, vol. 18, pp. 51–59, 2018.

[40] K. H. Maier-Hein, P. F. Neher, J.-C. Houde, M.-A. Côté, E. Garyfallidis, J. Zhong et al., “The challenge of mapping the human connectome based on diffusion tractography,” Nature communications, vol. 8, no. 1, pp. 1–13, 2017.

[41] L. J. O’Donnell, W. M. Wells, A. J. Golby, and C.-F. Westin, “Unbiased groupwise registration of white matter tractography,” in MICCAI, 2012, pp. 123–130.

[42] E. Garyfallidis, O. Ocegueda, D. Wassermann, and M. Descoteaux, “Robust and efficient linear registration of white-matter fascicles in the space of streamlines,” NeuroImage, vol. 117, pp. 124–140, 2015.

[43] E. Olivetti, N. Sharmin, and P. Avesani, “Alignment of tractograms as graph matching,” Front. Neuroscience, vol. 10, p. 554, 2016.

[44] I. Benou, R. Veksler, A. Friedman, and T. R. Raviv, “Combining white matter diffusion and geometry for tract-specific alignment and variability analysis,” NeuroImage, vol. 200, pp. 674–689, 2019.

[45] B. Q. Chandio and E. Garyfallidis, “StND: Streamline-based Non-rigid partial-Deformation Tractography Registration,” in Medical Imaging Meets NeurIPS, 2020.

[46] U. Ziyan, M. Sabuncu, L. J. O’Donnell, and C.-F. Westin, “Nonlinear registration of diffusion MR images based on fiber bundles,” in MICCAI, 2007, pp. 351–358.

[47] D. Wassermann, Y. Rathi, S. Bouix, M. Kubicki, R. Kikinis, M. Shenton et al., “White matter bundle registration and population analysis based on Gaussian processes,” in IPMI, 2011, pp. 320–332.

[48] S. Durrleman, P. Fillard, X. Pennec, A. Trouvé, and N. Ayache, “Registration, atlas estimation and variability analysis of white matter fiber bundles modeled as currents,” NeuroImage, vol. 55, no. 3, pp. 1073–1090, 2011.

[49] A. Sotiras, C. Davatzikos, and N. Paragios, “Deformable medical image registration: A survey,” IEEE TMI, vol. 32, no. 7, pp. 1153–1190, 2013.

[50] G. Haskins, U. Kruger, and P. Yan, “Deep learning in medical image registration: a survey,” Mach. Vis. Appl., vol. 31, no. 1, pp. 1–18, 2020.

[51] Y. Fu, Y. Lei, T. Wang, W. J. Curran, T. Liu, and X. Yang, “Deep learning in medical image registration: a review,” Phys. Med. Biol., vol. 65, no. 20, p. 20TR01, 2020.

[52] A. Sedghi, L. J. O’Donnell, T. Kapur, E. Learned-Miller, P. Mousavi, and W. M. Wells III, “Image registration: Maximum likelihood, minimum entropy and deep learning,” MedIA, vol. 69, p. 101939, 2021.

[53] I. Grigorescu, A. Uus, D. Christiaens, L. Cordero-Grande, J. Hutter, A. D. Edwards et al., “Diffusion tensor driven image registration: a deep learning approach,” in WBIR, 2020, pp. 131–140.

[54] G. Balakrishnan, A. Zhao, M. Sabuncu, J. Guttag, and A. Dalca, “Voxelmorph: a learning framework for deformable medical image registration,” IEEE TMI, vol. 38, no. 8, pp. 1788–1800, 2019.

[55] J. Wasserthal, P. F. Neher, D. Hirjak, and K. H. Maier-Hein, “Combined tract segmentation and orientation mapping for bundle-specific tractography,” MedIA, vol. 58, p. 101559, 2019.

[56] M. Jaderberg, K. Simonyan, A. Zisserman, and K. Kavukcuoglu, “Spatial transformer networks,” in NIPS, 2015, pp. 2017–2025.

[57] Z. Xu, J. Luo, J. Yan, R. Pulya, X. Li, W. Wells et al., “Adversarial uni-and multi-modal stream networks for multimodal image registration,” in MICCAI, 2020, pp. 222–232.

[58] B. B. Avants, C. L. Epstein, M. Grossman, and J. C. Gee, “Symmetric diffeomorphic image registration with cross-correlation: evaluating automated labeling of elderly and neurodegenerative brain,” MedIA, vol. 12, no. 1, pp. 26–41, 2008.

[59] Z. Gu, J. Cheng, H. Fu, K. Zhou, H. Hao, Y. Zhao et al., “Ce-net: Context encoder network for 2D medical image segmentation,” IEEE TMI, vol. 38, no. 10, pp. 2281–2292, 2019.

[60] Q. Lu, Y. Li, and C. Ye, “Volumetric white matter tract segmentation with nested self-supervised learning using sequential pretext tasks,” MedIA, vol. 72, p. 102094, 2021.

[61] M. H. Vu, G. Grimbergen, T. Nyholm, and T. Löfstedt, “Evaluation of multislice inputs to convolutional neural networks for medical image segmentation,” Medical Physics, 2020.

[62] A. Paszke, S. Gross, F. Massa, A. Lerer, J. Bradbury, G. Chanan et al., “Pytorch: An imperative style, high-performance deep learning library,” in NIPS, 2019, pp. 8026–8037.

[63] D. P. Kingma and J. Ba, “Adam: A method for stochastic optimization,” arXiv:1412.6980, 2014.

[64] I. Norton, W. Essayed, F. Zhang, S. Pujol, A. Yarmarkovich, A. Golby et al., “SlicerDMRI: open source diffusion MRI software for brain cancer research,” Cancer Res., vol. 77, pp. 101–103, 2017.

[65] F. Zhang, T. Noh, P. Juvekar, S. F. Frisken, L. Rigolo, I. Norton et al., “SlicerDMRI: Diffusion MRI and tractography research software for brain cancer surgery planning and visualization,” JCO Clin. Cancer Inform., vol. 4, pp. 299–309, 2020.

[66] M. Glasser, S. Sotiropoulos, A. Wilson, T. Coalson, B. Fischl et al., “The minimal preprocessing pipelines for the Human Connectome Project,” NeuroImage, vol. 80, pp. 105–124, 2013.

[67] B. Casey, T. Cannonier, M. I. Conley, A. O. Cohen, D. M. Barch, M. M. Heitzeg et al., “The adolescent brain cognitive development (ABCD) study: imaging acquisition across 21 sites,” Developmental cognitive neuroscience, vol. 32, pp. 43–54, 2018.

[68] K. Marek, D. Jennings, S. Lasch, A. Siderowf, C. Tanner, T. Simuni et al., “The Parkinson progression marker initiative (PPMI),” Progress in neurobiology, vol. 95, no. 4, pp. 629–635, 2011.

[69] D. J. Hagler Jr, S. Hatton, M. D. Cornejo, C. Makowski, D. A. Fair, A. S. Dick et al., “Image processing and analysis methods for the adolescent brain cognitive development study,” NeuroImage, vol. 202, p. 116091, 2019.

[70] F. Zhang, Y. Wu, I. Norton, L. Rigolo, Y. Rathi, N. Makris et al., “An anatomically curated fiber clustering white matter atlas for consistent white matter tract parcellation across the lifespan,” NeuroImage, vol. 179, pp. 429–447, 2018.

[71] J. D. Tournier, F. Calamante, and A. Connelly, “Improved probabilistic streamlines tractography by 2nd order integration over fibre orientation distributions,” in ISMRM, 2010, p. 1670.

[72] B. Jeurissen, J.-D. Tournier, T. Dhollander, A. Connelly, and J. Sijbers, “Multi-tissue constrained spherical deconvolution for improved analysis of multi-shell diffusion mri data,” NeuroImage, vol. 103, pp. 411–426, 2014.

[73] J. Wasserthal, P. Neher, and K. H. Maier-Hein, “Tractseg-fast and accurate white matter tract segmentation,” NeuroImage, vol. 183, pp. 239–253, 2018.

[74] F. Zhang, A. Breger, K. I. K. Cho, L. Ning, C.-F. Westin, L. J. O’Donnell et al., “Deep Learning Based Segmentation of Brain Tissue from Diffusion MRI,” NeuorImage, vol. 233, p. 117934, 2021.

[75] L. R. Dice, “Measures of the amount of ecologic association between species,” Ecology, vol. 26, no. 3, pp. 297–302, 1945.

[76] L. J. O’Donnell and C.-F. Westin, “Automatic tractography segmentation using a high-dimensional white matter atlas,” IEEE TMI, vol. 26, no. 11, pp. 1562–1575, 2007.

[77] A. Newell, “A tutorial on speech understanding systems,” Speech recognition, pp. 4–54, 1975.

[78] J. Cohen, Statistical power analysis for the behavioral sciences. Academic press, 2013.

[79] A. Klein, J. Andersson, B. A. Ardekani, J. Ashburner, B. Avants, M.-C. Chiang et al., “Evaluation of 14 nonlinear deformation algorithms applied to human brain MRI registration,” NeuroImage, vol. 46, no. 3, pp. 786–802, 2009.

[80] B. B. Avants, N. Tustison, and G. Song, “Advanced normalization tools (ANTS),” Insight J, vol. 2, no. 365, pp. 1–35, 2009.

[81] Y. Wang, A. Gupta, Z. Liu, H. Zhang, M. L. Escolar, J. H. Gilmore et al., “DTI registration in atlas based fiber analysis of infantile Krabbe disease,” NeuroImage, vol. 55, no. 4, pp. 1577–1586, 2011.

[82] M. Jenkinson, C. F. Beckmann, T. E. Behrens, M. W. Woolrich, and S. M. Smith, “Fsl,” NeuroImage, vol. 62, no. 2, pp. 782–790, 2012.

[83] A. D. Leow, I. Yanovsky, M.-C. Chiang, A. D. Lee, A. D. Klunder, A. Lu et al., “Statistical properties of jacobian maps and the realization of unbiased large-deformation nonlinear image registration,” IEEE TMI, vol. 26, no. 6, pp. 822–832, 2007.

[84] M. Mee, K. Stewart, M. Lathouras, H. Truong, and C. Hargrave, “Evaluation of a deformable image registration quality assurance tool for head and neck cancer patients,” J. Med. Rad. Sci., vol. 67, no. 4, pp. 284–293, 2020.

[85] J. Ashburner, “A fast diffeomorphic image registration algorithm,” NeuroImage, vol. 38, no. 1, pp. 95–113, 2007.

[86] D. Rueckert, L. I. Sonoda, C. Hayes, D. L. Hill, M. O. Leach, and D. J. Hawkes, “Nonrigid registration using free-form deformations: application to breast MR images,” IEEE TMI, vol. 18, no. 8, pp. 712–721, 1999.

[87] Y. Hu, M. Modat, E. Gibson, W. Li, N. Ghavami, E. Bonmati et al., “Weakly-supervised convolutional neural networks for multimodal image registration,” MedIA, vol. 49, pp. 1–13, 2018.

[88] W. Zhu, A. Myronenko, Z. Xu, W. Li, H. Roth, Y. Huang et al., “Neurreg: Neural registration and its application to image segmentation,” in CVPR, 2020, pp. 3617–3626.

[89] A. Sedghi, T. Kapur, J. Luo, P. Mousavi, and W. M. Wells, “Probabilistic image registration via deep multi-class classification: characterizing uncertainty,” in MICCAI-UNSURE, 2019, pp. 12–22.

[90] S. Pajevic and C. Pierpaoli, “Color schemes to represent the orientation of anisotropic tissues from diffusion tensor data: application to white matter fiber tract mapping in the human brain,” Mag. Res. Med., vol. 42, no. 3, pp. 526–540, 1999.

[91] R. Polikar, “Ensemble learning,” in Ensemble machine learning, 2012, pp. 1–34.

[92] C. Zhang and Y. Ma, Ensemble machine learning: methods and applications. Springer, 2012.

[93] B. Lakshminarayanan, A. Pritzel, and C. Blundell, “Simple and scalable predictive uncertainty estimation using deep ensembles,” in NIPS, 2017, p. 6405–6416.

[94] O. Sagi and L. Rokach, “Ensemble learning: A survey,” Wiley Interdisciplinary Reviews: Data Mining and Knowledge Discovery, vol. 8, no. 4, p. e1249, 2018.

[95] Y. Cao, T. A. Geddes, J. Y. H. Yang, and P. Yang, “Ensemble deep learning in bioinformatics,” Nature Machine Intelligence, vol. 2, no. 9, pp. 500–508, 2020.

[96] C. Thomas, Q. Y. Frank, M. O. Irfanoglu, P. Modi, K. S. Saleem, D. A. Leopold et al., “Anatomical accuracy of brain connections derived from diffusion MRI tractography is inherently limited,” PNAS, vol. 111, no. 46, pp. 16 574–16 579, 2014.

[97] V. Nath, S. K. Pathak, K. G. Schilling, W. Schneider, and B. A. Landman, “Deep learning estimation of multi-tissue constrained spherical deconvolution with limited single shell DW-MRI,” in SPIE, vol. 11313, 2020, p. 113130S.

