## supplementary for "Deep Diffusion MRI Registration (DDMReg): A Deep Learning Method for Diffusion MRI Registration"

| Supplementary Table S1. List of TOMs [47] |
|---|
| 1: AF - Arcuate fascicle<br>2: ATR - Anterior thalamic radiation<br>3: CA - Commissure anterior<br>4: CC_1 - Rostrum<br>5: CC_2 - Genu<br>6: CC_3 - Rostral body (Premotor)<br>7: CC_4 - Anterior midbody (Primary motor)<br>8: CC_5 - Posterior midbody (Primary somatosensory)<br>9: CC_6 - Isthmus<br>10: CC_7 - Splenium<br>11: CG - Cingulum left<br>12: CST - Corticospinal tract<br>13: MLF - Middle longitudinal fascicle<br>14: FPT - Fronto-pontine tract<br>15: FX - Fornix<br>16: ICP - Inferior cerebellar peduncle<br>17: IFO - Inferior occipito-frontal fascicle<br>18: ILF - Inferior longitudinal fascicle<br>19: MCP - Middle cerebellar peduncle<br>20: OR - Optic radiation<br>21: POPT - Parieto-occipital pontine<br>22: SCP - Superior cerebellar peduncle<br>23: SLF_I - Superior longitudinal fascicle I<br>24: SLF_II - Superior longitudinal fascicle II<br>25: SLF_III - Superior longitudinal fascicle III<br>26: STR - Superior thalamic radiation<br>27: T_PREF - Thalamo-prefrontal<br>28: T_PREM - Thalamo-premotor<br>29: T_PREC - Thalamo-precentral<br>30: T_POSTC - Thalamo-postcentral<br>31: T_PAR - Thalamo-parietal<br>32: T_OCC - Thalamo-occipital<br>33: ST_FO - Striato-fronto-orbital<br>34: ST_PREF - Striato-prefrontal<br>35: ST_PREM - Striato-premotor<br>36: ST_PREC - Striato-precentral<br>37: ST_POSTC - Striato-postcentral<br>38: ST_PAR - Striato-parietal<br>39: ST_OCC - Striato-occipital<br>40: UF - Uncinate fascicle |

Supplementary Table S2. Statistical comparison across the $Reg_{FA}$, $Reg_{FA+TOM}$, and $Reg_{proposed}$ methods.

| Evaluation metric | Tract Dice score | | Tissue Dice score | | Tract Distance | |
|---|---|---|---|---|---|---|
| Three-method comparison (one-way repeated measures ANOVA) | $F = 116.57$; $p < 0.001$ * | | $F = 183.33$; $p < 0.001$ * | | $F = 41.15$; $p < 0.001$ * | |
| Pairwise comparison (paired t-test) | p-value | Cohen's d | p-value | Cohen's d | p-value | Cohen's d |
| $Reg_{proposed}$ vs $Reg_{FA+TOM}$ | < 0.001 * | 1.72 | < 0.001 * | 1.12 | < 0.001 * | 0.72 |
| $Reg_{proposed}$ vs $Reg_{FA}$ | < 0.001 * | 3.00 | < 0.001 * | 4.72 | < 0.001 * | 1.37 |
| $Reg_{FA+TOM}$ vs $Reg_{FA}$ | < 0.001 * | 2.61 | < 0.001 * | 5.40 | < 0.001 * | 1.53 |

Supplementary Table S3. Statistical comparison across the SyN, DTI-TK, MRRegister, VoxelMorph and DDMReg methods.

| Evaluation metric | Tract Dice score | | Tissue Dice score | | Tract Distance | |
|---|---|---|---|---|---|---|
| Five-method comparison (one-way repeated measures ANOVA) | F = 506.47; p < 0.001 * | | F = 797.48; p < 0.001 * | | F = 521.44; p < 0.001 * | |
| Pairwise comparison (paired t-test) | p-value | Cohen's d | p-value | Cohen's d | p-value | Cohen's d |
| DDMReg vs VoxelMorph | < 0.001 * | 3.19 | < 0.001 * | 3.64 | < 0.001 * | 2.40 |
| DDMReg vs MRRegister | < 0.001 * | 0.93 | < 0.001 * | 1.31 | < 0.001 * | 0.84 |
| DDMReg vs DTI-TK | < 0.001 * | 0.56 | < 0.001 * | 1.16 | < 0.001 * | 1.60 |
| DDMReg vs SyN | < 0.001 * | 2.00 | < 0.001 * | 2.33 | < 0.001 * | 1.63 |
| VoxelMorph vs MRRegister | < 0.001 * | 1.02 | < 0.001 * | 1.01 | < 0.001 * | 0.87 |
| VoxelMorph vs DTI-TK | < 0.001 * | 0.59 | < 0.001 * | 0.84 | < 0.001 * | 1.20 |
| VoxelMorph vs SyN | < 0.001 * | 1.17 | < 0.001 * | 1.21 | < 0.001 * | 1.15 |
| MRRegister vs DTI-TK | < 0.001 * | 0.91 | < 0.001 * | 0.93 | < 0.001 * | 0.90 |
| MRRegister vs SyN | < 0.001 * | 1.04 | < 0.001 * | 0.94 | < 0.001 * | 0.82 |
| DTI-TK vs SyN | < 0.001 * | 0.43 | < 0.001 * | 0.82 | < 0.001 * | 1.04 |

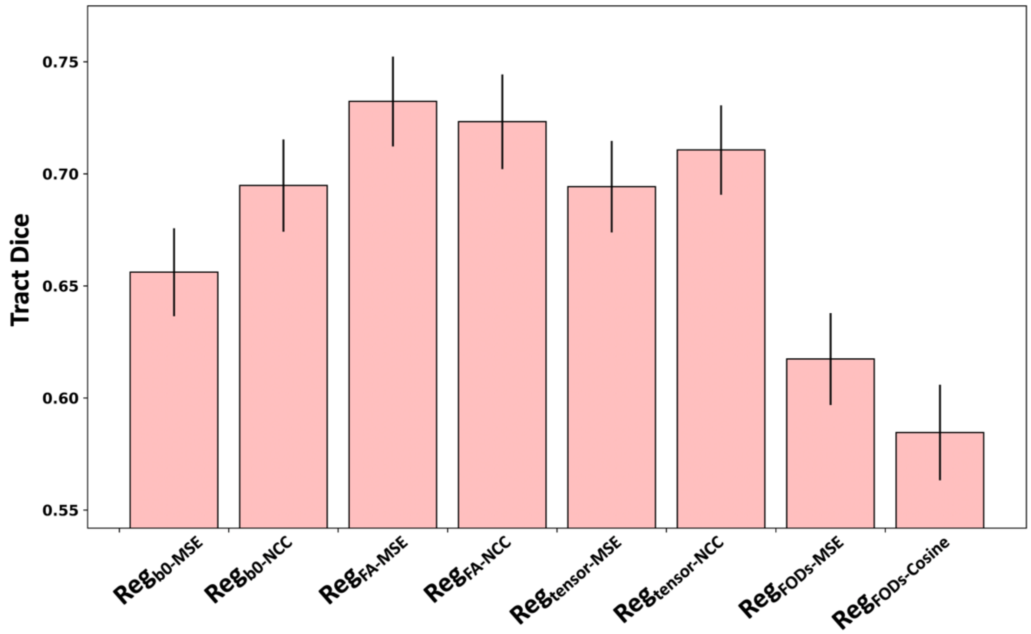


Supplementary Fig. S1. The performance of whole-brain registration subnetwork (i.e., the U-net architecture as proposed in VoxelMorph) when using different inputs and loss functions.

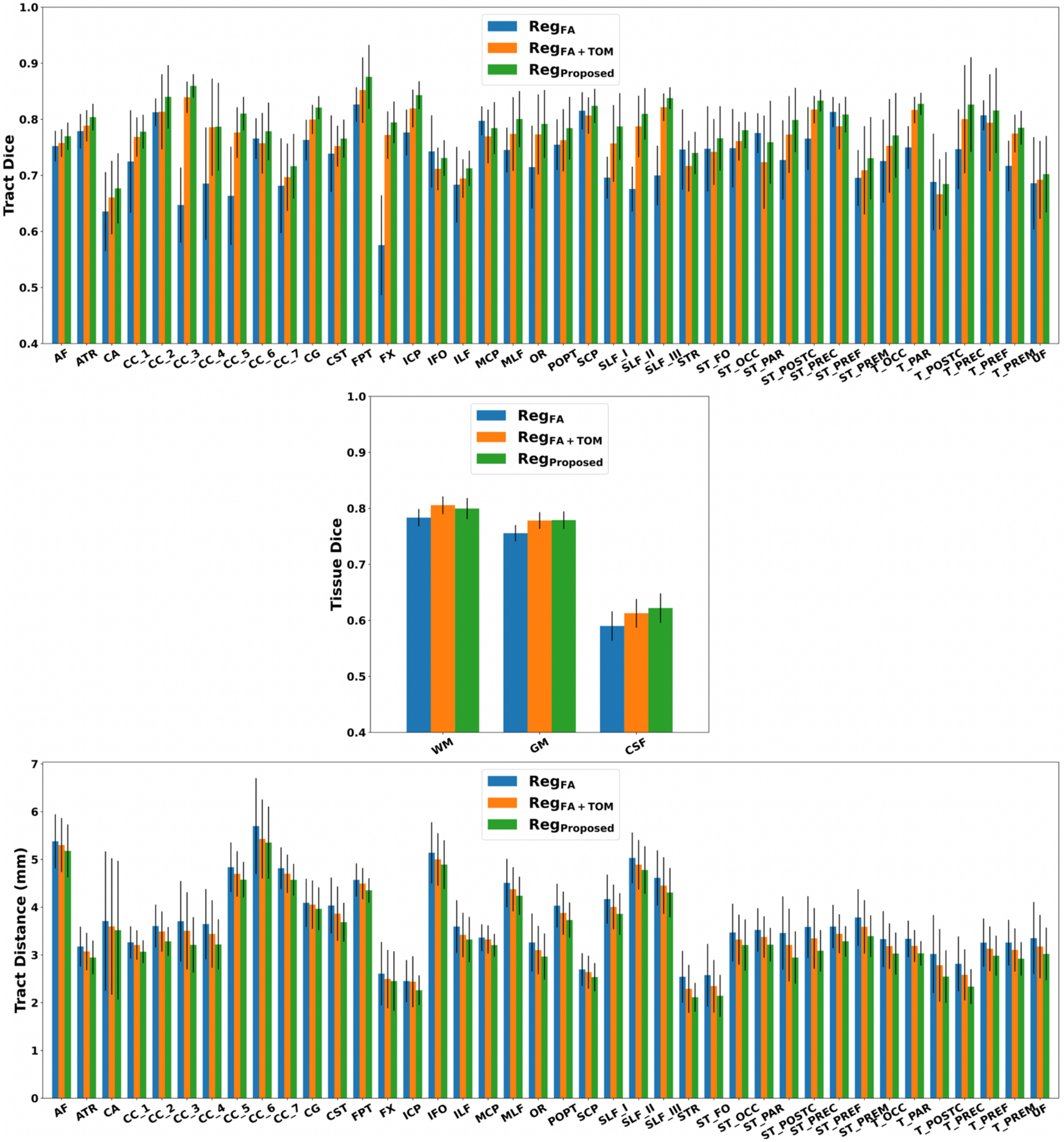


Supplementary Fig. S2. Individual metric values of the Reg$_{FA}$, Reg$_{FA+TOM}$, and Reg$_{proposed}$ methods.

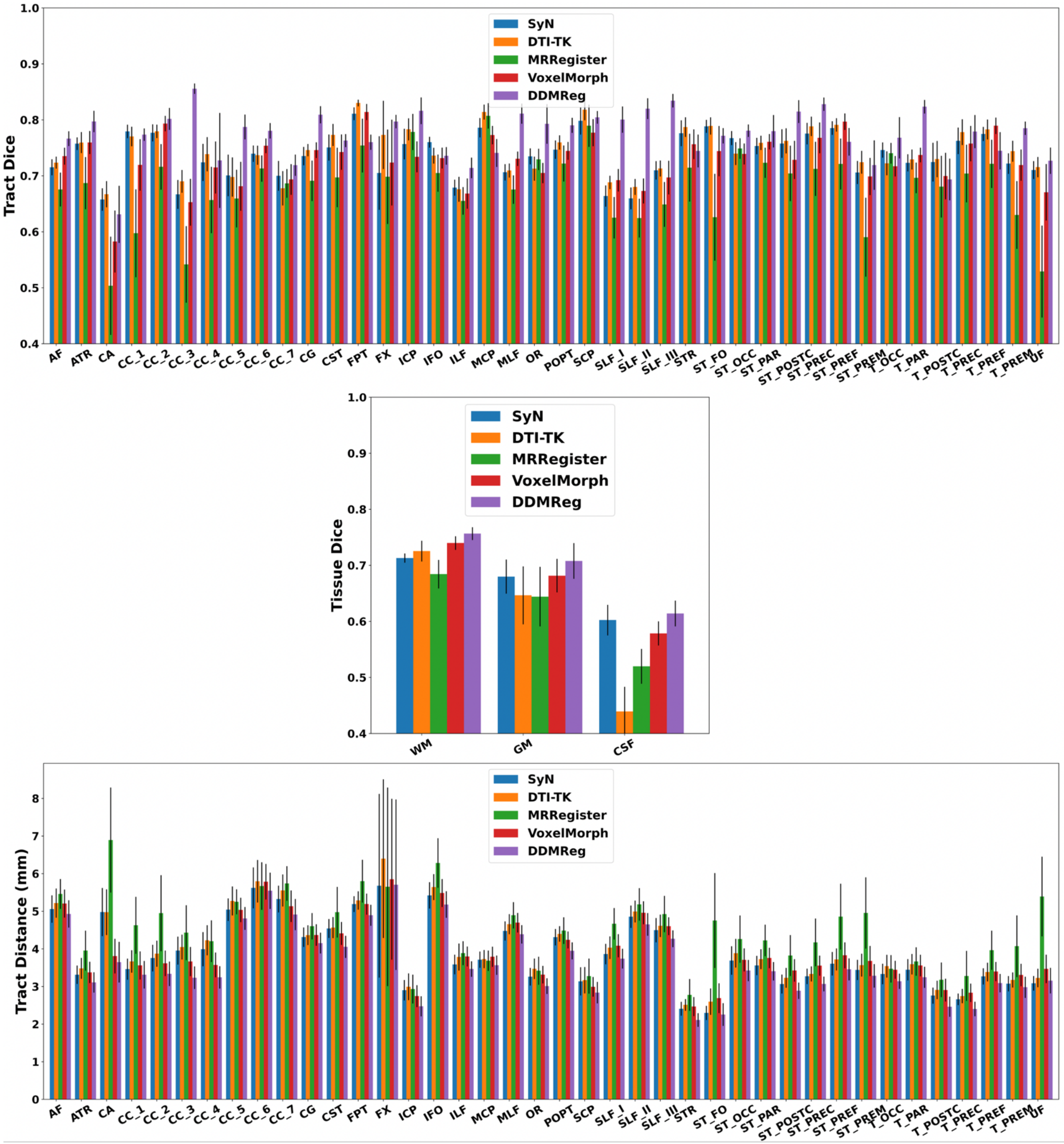


Supplementary Fig. S3. Individual metric values of the SyN, DTI-TK, MRRegister, VoxelMorph and DDMReg methods.

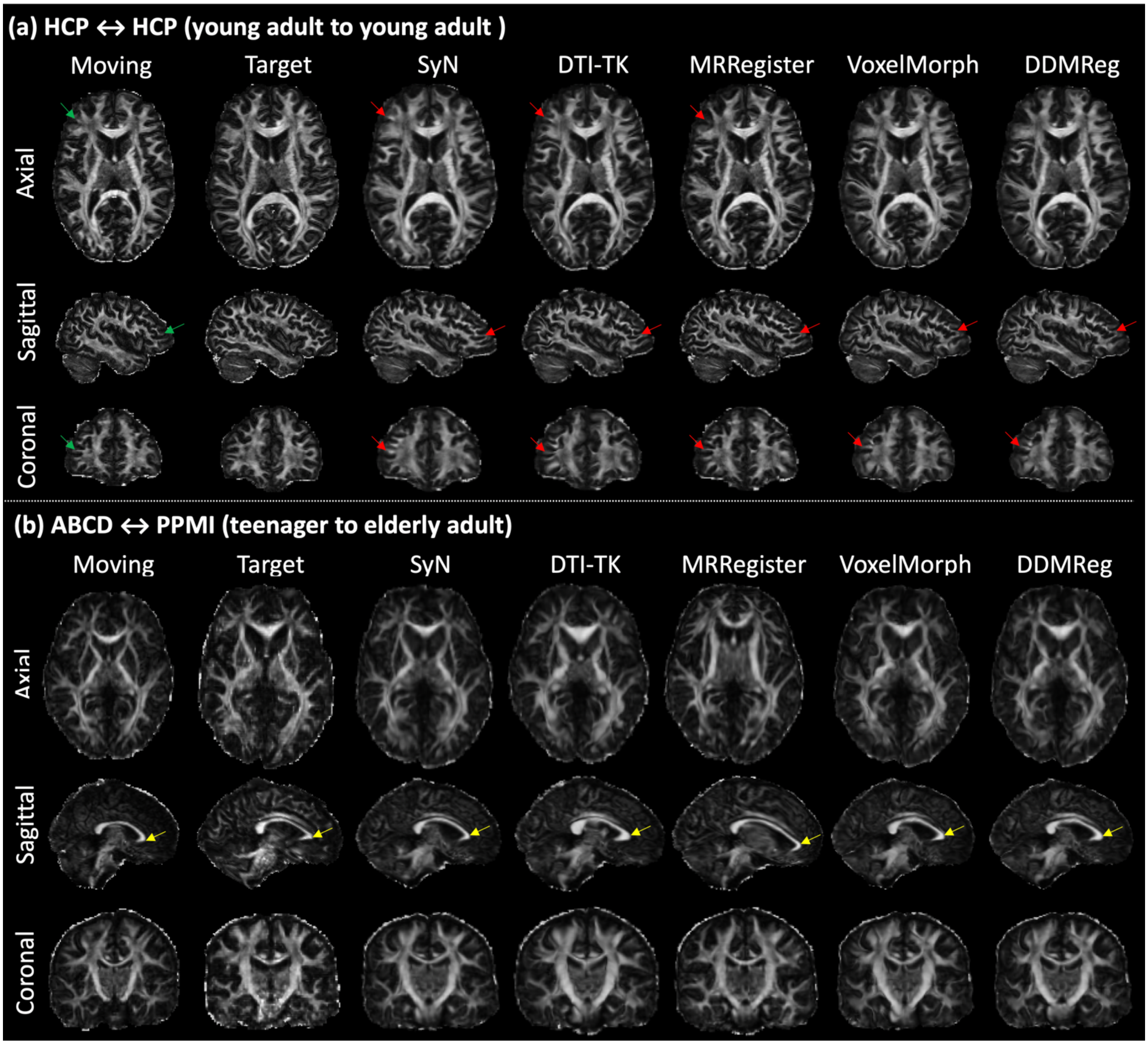


Supplementary Fig. S4: Visualization of registration results for the state-of-the-art and DDMReg methods. This figure provides additional visualization of the results in Fig. 6 of the main paper by adding the sagittal and coronal views. In subfigure (a), for each compared method, red arrows are added in the sagittal and coronal views to show the registered white matter structure that corresponds to the structure (indicated using blue arrows) in the moving image. In subfigure (b), for each compared method, a yellow arrow is added in the sagittal view to indicate the anterior part of the corpus callosum, providing additional visualization showing the differences of the registration across methods.

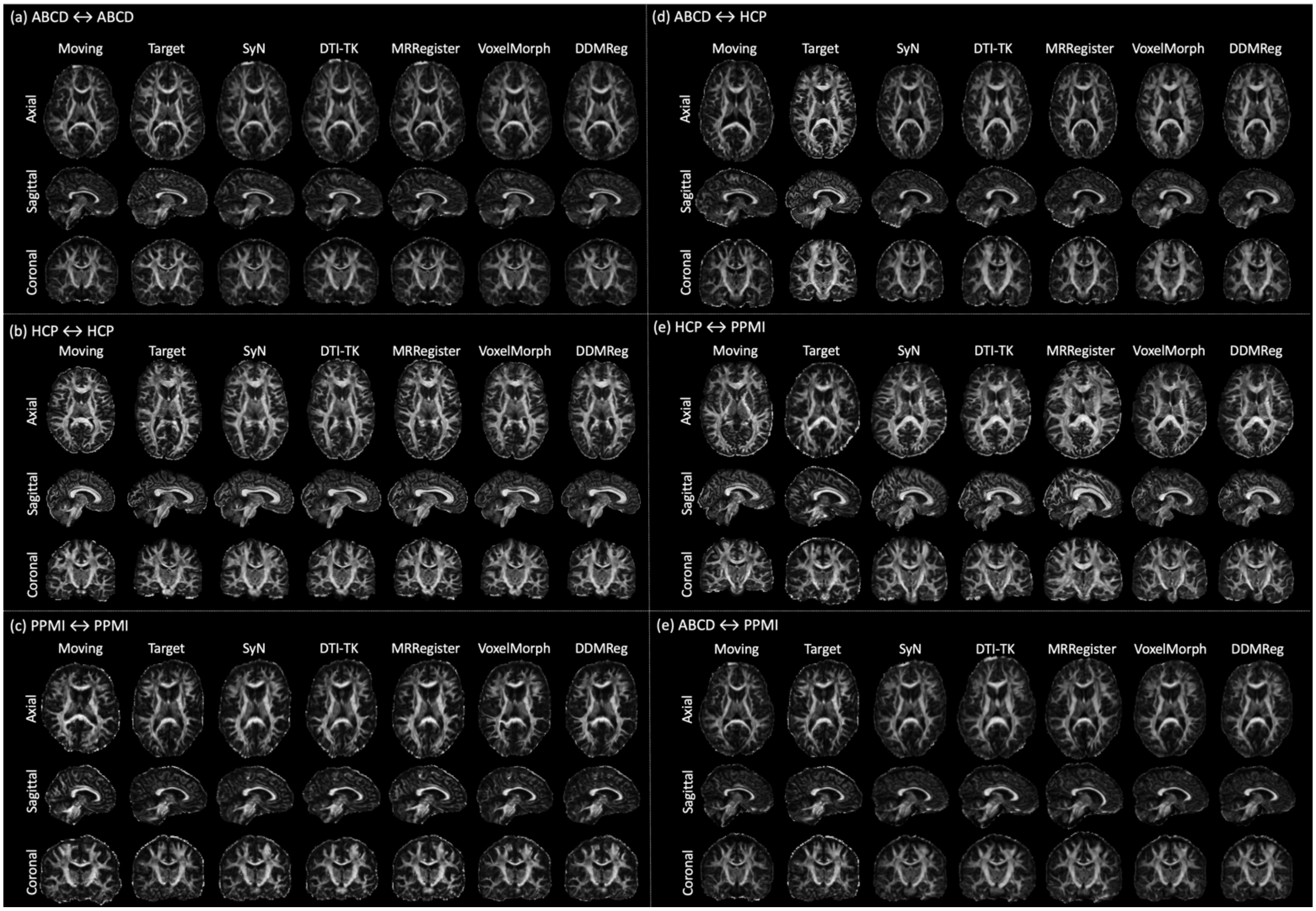


Supplementary Fig. S5: Visualization of registration results for the state-of-the-art and DDMReg methods. Six examples are provided, where subfigures (a) to (c) are intra-population registration examples and (d) to (f) are inter-population registration examples.

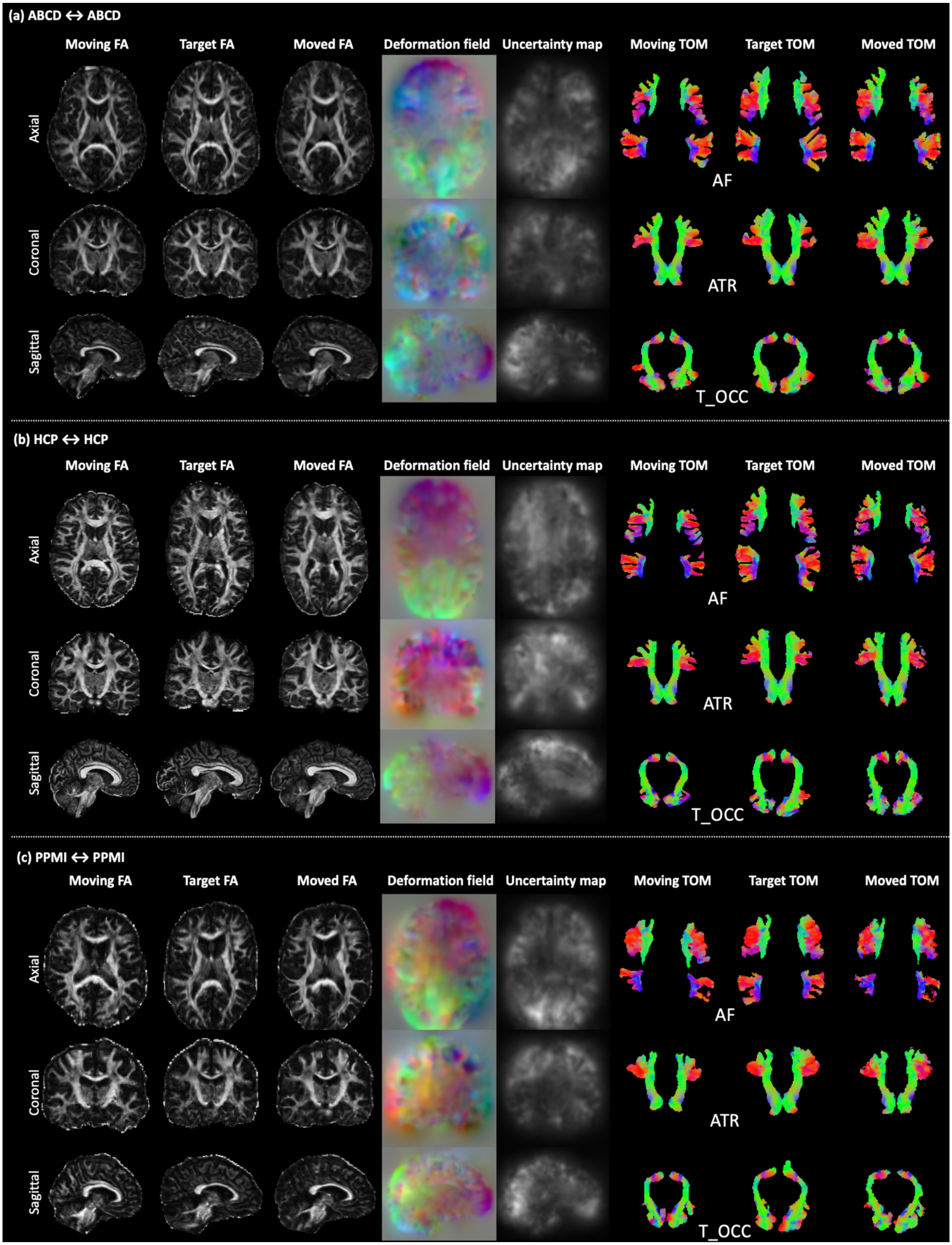


Supplementary Fig. S6. Visualization of registration results of three additional example intra-population image pairs using DDMReg. Images from axial, coronal, and sagittal views are provided for FA, the estimated deformation field, and the uncertainty map. TOMs of three example tracts (AF, ATR, and T_OCC) are provided from the axial view.

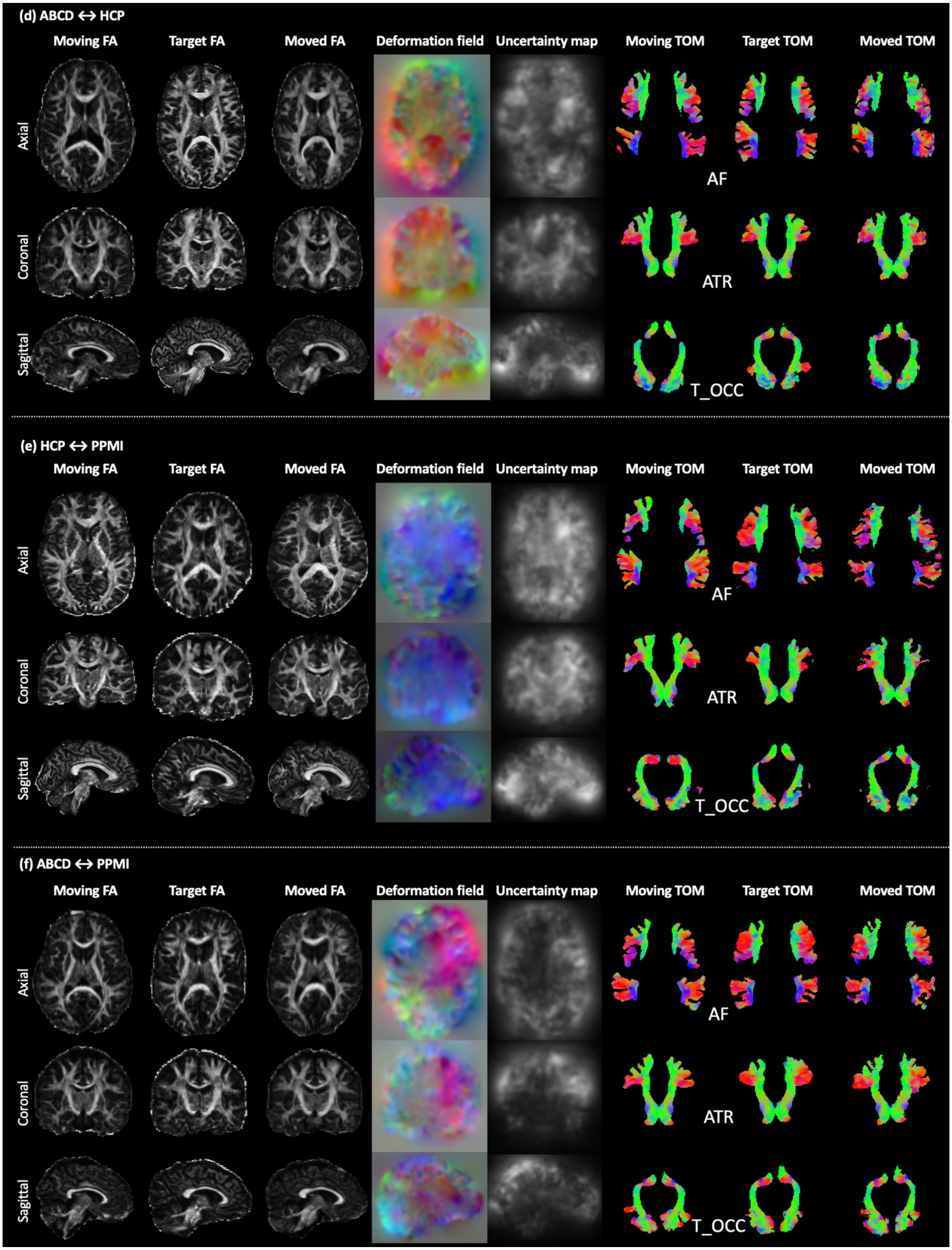


Supplementary Fig. 7. Visualization of registration results of three additional example inter-population image pairs using DDMReg. Images from axial, coronal, and sagittal views are provided for FA, the estimated deformation field, and the uncertainty map. TOMs of three example tracts (AF, ATR, and T_OCC) are provided from the axial view.
